# Modeling gene expression evolution with EvoGeneX uncovers differences in evolution of species, organs and sexes

**DOI:** 10.1101/2020.01.06.895615

**Authors:** Soumitra Pal, Brian Oliver, Teresa M. Przytycka

## Abstract

While DNA sequence evolution has been well studied, the expression of genes is also subject to evolution, yet the evolution of gene expression is currently not well understood. Recently, new tissue/organ-specific gene expression datasets spanning several organisms across the tree of life, have become available, providing the opportunity to study gene expression evolution in more detail. While a theoretical model to study the evolution of continuous traits exists, in practice computational methods often cannot confidently distinguish between alternative evolutionary scenarios. This lack of power has been attributed to modest numbers of species considered in these studies.

We hypothesised that biological replicates can be used to increase predictive power of these models. With this in mind, we introduce EvoGeneX, a computationally efficient method to uncover the mode of gene expression evolution based on the Ornstein-Uhlenbeck process. Importantly, in addition to modelling expression variations between species, EvoGeneX models within-species variation.

Furthermore, to facilitate comparative analysis of gene expression evolution, we introduce a formal approach, based on Michaelis-Menten equation, to measure the dynamics of evolutionary divergence of a group of genes in terms of group’s asymptotic divergence level and rate.

Finally, we used these tools to preform the first analysis the evolution of gene expression across different body parts, species, and sexes of the *Drosophila* genus. Our analysis revealed that neutral expression evolution can be confidently rejected in favor of purifying selection in nearly half of the genes. In addition, we quantified differences in the evolutionary dynamics of male and female gonads and uncovered interesting examples of adaptive gene expression evolution.

## 1 Introduction

Studies of species evolution typically focus on the evolution of DNA sequence. However, in complex multi-cellular organisms all cells utilize the same genetic information, yet they show remarkable phenotypic differences arising from distinct transcriptional programs executed in different organs, tissues, and cells. While evolution ultimately acts at the level of the successful reproduction of individuals, divergent transcriptional programs within the individual are subject to evolutionary adaptation constraint and drift. Evolutionary analysis of gene expression can thus shed light on the evolutionary processes in ways that cannot be achieved by the analyses of sequence alone, and will provide a richer understanding of evolution.

Currently, our understanding of the interplay between species evolution and tissue-specific expression evolution is still limited. The results of initial analyses of gene expression evolution have not provided a clear picture of the dominant forces. For example, early studies of the evolution of primate gene expression suggested that expression evolution is largely consistent with the neutral theory of evolution (Khaitovich et al. 2004, 2005) while subsequent analyses uncovered signatures of positive selection in brain (Khaitovich et al. 2006b) and testis (Khaitovich et al. 2006a).

In the past decade, several gene expression datasets encompassing a larger number of species have been collected and analysed [mammals: (Brawand et al. 2011; Romero et al. 2012; J. Chen et al. 2019), vertebrates: (Chan et al. 2009), primates: (Blekhman et al. 2008)]. Studies in mammals indicated that, when restricted to individual tissues or organs, gene expression evolution tends to follow the species tree fairly closely (J. Chen et al. 2019). However multi-tissue samples generally clustered first by tissues rather than by species (or study) (Chan et al. 2009; Brawand et al. 2011; Breschi et al. 2016; J. Chen et al. 2019), suggesting strong evolutionary constraints on tissue/organ specific gene expression. Yet, a recent study identified sets of genes that cluster first by species (Breschi et al. 2016). Finally, sex-biased expression also imposes constraints on evolution (Meiklejohn et al. 2003). However the nature of constraints imposed by combination of tissue and sex specificity is less clear. In addition, tissue and sex specific studies of gene expression adaptation are very limited.

Recently, we collected a large dataset of body part and sex specific gene expression focusing on the *Drosophila* phylogeny (Yang et al. 2018). We use this data along with new computational tools developed in this study, to examine modes of gene expression evolution of *Drosophila* genus in the context of body parts, sex, and species groups.

Following the pioneering work of Felsenstein (1973), neutral evolution of continuous traits, such as gene expression, is formally modelled by Brownian Motion (BM). Along a similar line of thought, Lande (1976) pioneered the use of the Ornstein-Uhlenbeck (OU) process to model evolutionary constraints and adaptive evolution of continuous traits. The OU process is stochastic and extends the BM model by adding an “attraction force” towards an optimum value. Combining the OU process on gene expression with the information about the underlining evolutionary tree (obtained, for example, from sequence analysis) allows modeling the differences in evolution along individual branches and thus helps uncover branch specific adaptation (Hansen 1997; Butler and A. A. King 2004; Brawand et al. 2011; J. Chen et al. 2019).

Phylogenetically diverse gene expression datasets that span many organisms across the tree of life and include expression data on multiple tissues/organs have challenged us to put these theoretical models into practical use. For example, several recent works used stochastic models to study evolution of gene expression (Bedford and Hartl 2009; Kalinka et al. 2010; Nourmohammad et al. 2017; Brawand et al. 2011; J. Chen et al. 2019). Unfortunately, these formal stochastic models for continuously varying phenotypic traits usually do not account for within-species variation. Yet, the gene expression differs even between genetically identical multi-cellular organisms, such as *Drosophila*, where there are significant individual to individual variation (Lee et al. 2018; Lin et al. 2016). It has been long appreciated that ignoring within species variation might bias the results in comparative data analysis (Joseph Felsenstein 2008; Ives et al. 2007; Rohlfs et al. 2014). Within-species variation was considered in the initial analysis of mammalian gene expression (Brawand et al. 2011). Unfortunately, this study considered only constrained and adaptive evolution models ignoring the possibility of neutral evolution, even though neutral evolution predominates both within and between species variation (Kimura 1968; J. L. King and Jukes 1969). In addition, no attention to computational efficiency of the approach or a demonstration that accounting for within species variation impacted the results was given. Indeed, a subsequent analysis of extended mammalian phylogeny showed that neutral evolution of gene expression cannot be rejected for a large fraction of mammalian genes (J. Chen et al. 2019). However, this last study did not consider the within-species variation which is essential for understanding the meaning of variation between species.

Our contributions here are two-fold. First, given the above conflicting results and having in mind the need to facilitate future analyses of gene expression evolution, we developed a rigorous and computationally efficient approach, EvoGeneX, to model gene expression evolution for a given set of species under the assumption that the expression data for each species includes biological replicates. Next, we used EvoGeneX along with other methods we developed in this study to perform an extensive analysis of tissue and sex biased gene expression evolution in *Drosophila*. Mathematically, EvoGeneX follows the broadly accepted stochastic model of expression evolution and additionally supports the rigorous inclusion of replicates. Specifically, given the phylogenetic tree of a set of species (derived, for example, based on sequence evolution) and biologically replicated measurements of expression of a gene for all extant species in the tree, EvoGeneX, infers the most likely mode of expression evolution for the expression of that gene. We show that by modelling the within-species diversity EvoGeneX improves over the computational method OUCH (A. A. King and Butler 2009) that is currently accepted as the basic framework for gene expression evolution (J. Chen et al. 2019). EvoGeneX is developed under the same basic model as OUCH, but, by incorporating replicates to model within species variation, EvoGeneX has increased precision in detecting constrained and adaptive evolution. At the same time, thanks to our improved Maximum Likelihood estimation, EvoGeneX is very efficient despite the increased size of analysed data and increased number of parameters to be optimised.

We applied EvoGeneX to analyse our expression dataset encompassing 5 different body parts (head, gonads, thorax, viscera, and abdomen), from carefully selected representatives of the *Drosophila* genus (both male and female). For each of the 10 total sample types gene expression was measured by RNAseq in 4 biological replicates (Yang et al. 2018). The genus *Drosophila* is particularly well suited for studying gene expression evolution. The last common ancestor of the genus is assumed to date to the Cretaceous period about 112 *±* 28 million years ago (Wheat and Wahlberg 2013). The *Drosophila* species occupy diverse geographic locations and ecological niches (Morales-Hojas and Vieira 2012). Compared to the previous studies in mammals (Brawand et al. 2011; J. Chen et al. 2019) and vertebrates (Chan et al. 2009), the morphology of the *Drosophila* species is similar while at the same time the evolutionary distances are significantly larger relative to mammals or even vertebrates typically included in such studies. This makes *Drosophila* an ideal phylogeny to study the interplay among different modes of gene expression evolution.

Our analysis demonstrated that, in *Drosophila*, constrained evolution of gene expression is more common than neutral evolution, however, neutral evolution of gene expression cannot be rejected in a very large fraction of the genes. We found that expression of many genes is evolutionary constrained in multiple body parts. However, not all genes follow this pattern and have different evolutionary trends depending on where they are expressed. To gain further understanding of variations in evolutionary dynamics between organs and sexes, we introduced an approach based on Michaelis-Menten kinetics (Michaelis and Menten 1913) from enzyme kinetics theory. Our approach revealed striking differences in evolutionary dynamics of gene expression in male and female gonads, for example. We provide a specific analysis of adaptive expression evolution at the subgenus level. EvoGeneX revealed compelling examples of adaptive evolution of gene expression in the *Sophophora*/*Drosophila* branches of the phylogeny.

## 2 Results and Discussion

### 2.1 EvoGeneX: A fast and rigorous replica-aware approach to model gene expression evolution

Given a gene with one-to-one orthology relation across a set of species, the species tree, and expression values of the gene measured in multiple biological replicates for each species, our goal is to uncover the most likely scenario for the gene expression evolution. The three basic modes of evolution considered in this study are: (i) neutral evolution, (ii) constrained evolution (purifying/stabilising selection) where the evolution of gene expression is biased against divergence from some optimum value, and (iii) adaptive evolution where the expression in different groups of species is biased towards different optimum values.

Following standard practice, we assume that the evolution of gene expression follows a stochastic process. In particular, we assume that neutral evolution follows Brownian Motion (BM) – a special case of the more general mean-reverting Ornstein-Uhlenbeck (OU) process – the broadly accepted model for the evolution of continuous traits such as gene expression (Hansen 1997; Butler and A. A. King 2004; A. A. King and Butler 2009; Bedford and Hartl 2009; Brawand et al. 2011; J. Chen et al. 2019). This model assumes that gene expression follows a stochastic process that is “attracted” towards some optimum value. The strength of the bias is modelled as a constant *α*. The optimum value to which the process is attracted is allowed to change over the evolutionary time, reflecting changes in environmental or other constraints acting on the trait.

Assuming neutral evolution as a null model, our goal is to use a hypothesis testing framework to uncover alternative modes of gene expression evolution. Towards this end we developed EvoGeneX, a mathematical framework to estimate the parameters of the process given the data. To account for the within-species variability, we assume that for each species *i*, there are several observations reflecting the within-species variability of the trait of interest (here expression of a specific gene). Denoting the value of the *k*th observation of the trait for species *i* by *y*_*i,k*_ we set:

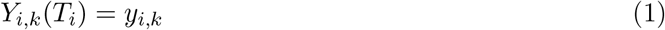

where *T*_*i*_ is the evolutionary time from the least common ancestor of all species in this tree to species *i*. Furthermore, for *t ≤ T*_*i*_ we define:

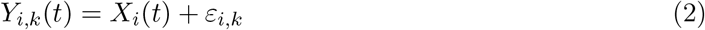

where *X*_*i*_(*t*) follows the OU process:

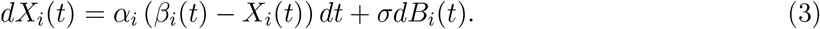

In equation (3) the term *σdB*_*i*_(*t*) models the increments of standard Brownian Motion (BM), *β*_*i*_(*t*) is the optimum trait value for species *i* at time *t*, and *α*_*i*_ is the strength of the attraction towards the optimum value. In addition, *β*_*i*_(*t*) is assumed to change in discrete steps corresponding to the speciation events (internal nodes of the evolutionary tree) (Hansen 1997; Butler and A. A. King 2004; J. Chen et al. 2019). Finally, *ε*_*i,k*_ ∼ *N* (0, *γσ*^2^) is an identically distributed Gaussian variable with mean 0 and variance *γσ*^2^ that models the within-species variability. We assume that within-species variance is smaller than evolutionary variance, hence the factor *γ* is assumed to be less than 1.

Given this general framework, we use hypothesis testing (Fig. 1d) to differentiate among the following evolutionary models: (i) **neutral evolution:** *α* = 0, (ii) **constrained evolution:** *α >* 0 and one common optimum *β*_*i*_(*t*) = *θ*_0_ for all *i* and *t*, and (iii) **adaptive evolution (evolutionary shift):** *α >* 0 and two different optima *θ*_0_, *θ*_1_ representing two regimes with optimal *β* values *θ*_1_ in a specific subtree and *θ*_0_ in the rest of the evolutionary tree. In all cases appropriate *θ* values together with *α, σ, γ* must be estimated from the data (i.e. values *y*_*i,k*_). Towards this end, in the Methods section, we describe our efficient method to compute Maximum Likelihood estimates of model parameters.

**Figure 1:**
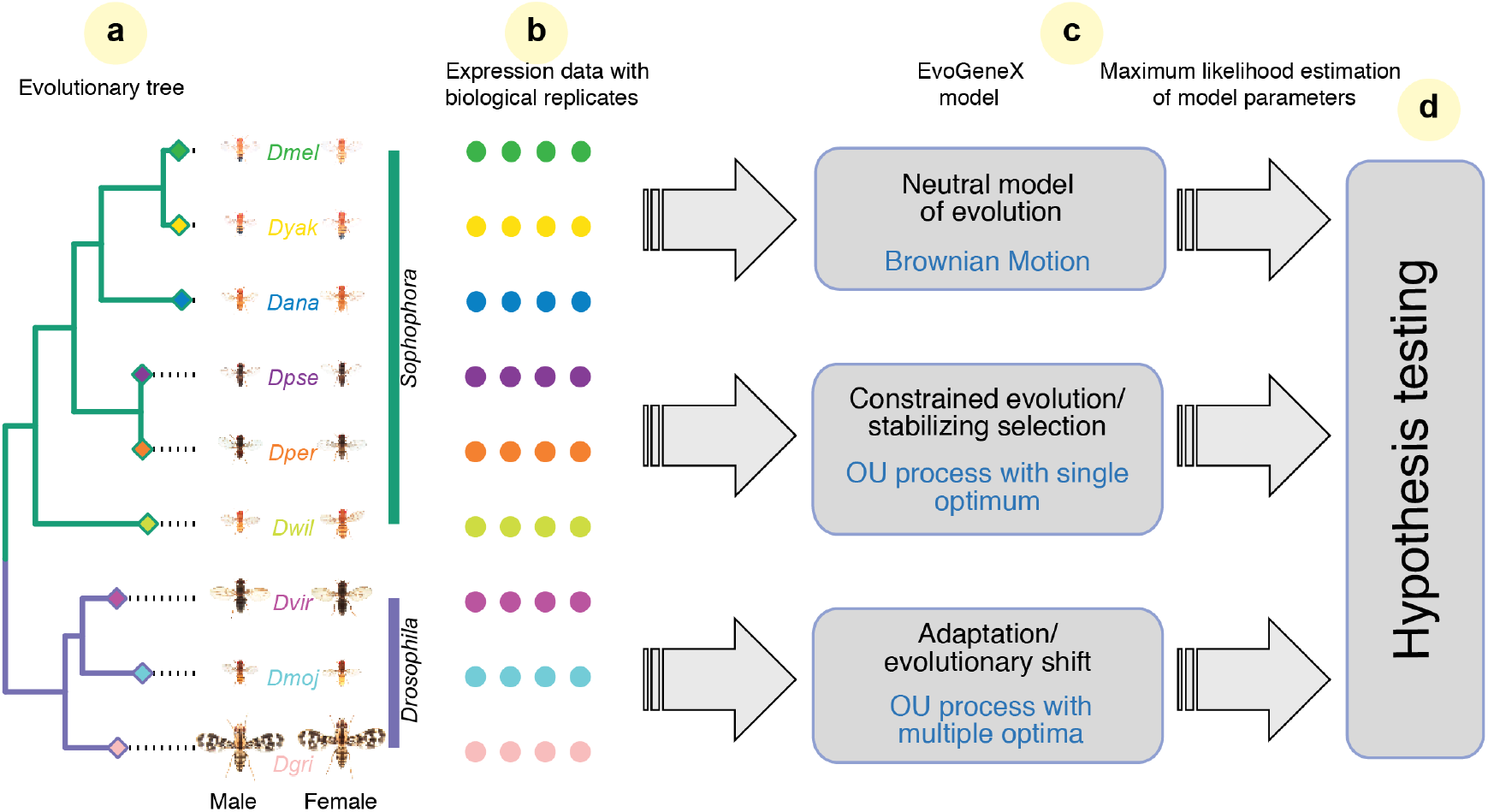
Workflow diagram. EvoGeneX takes as input an evolutionary tree (including evolutionary distances as branch lengths) (**a**) and normalised expression values across biological replicates and species (**b**). It models evolutionary scenarios as stochastic processes as described in the text and uses maximum likelihood approach to fit the parameters of the model to the data (**c**). For the adaptive model, the proposed adaptation regimes (here *Sophophora* and *Drosophila* subgroups) are required as part of the input. Finally, model selection is performed using hypothesis testing (**d**). Photos of flies from *Drosophila* species are courtesy of Nicolas Gompel.

### 2.2 EvoGeneX outperforms the previous leading approach

We use simulations to compare EvoGeneX with OUCH (A. A. King and Butler 2009) – currently the most popular software implementing an OU-based approach (Hansen 1997; Butler and A. A. King 2004; Bedford and Hartl 2009; J. Chen et al. 2019). We simulated expression values for 1000 genes using the stochastic differential equations (2) and (3) governing our OU-based model of gene expression evolution on phylogenetic trees, for each setting of the parameters *α, σ, β* and *γ* subject to biologically realistic constraints discussed below. For realistic evaluation of the effectiveness of our approach, we use for the simulation the same sequence based phylogenetic tree (Z.-X. Chen et al. 2014), that we use for the real data. To match our real data, we simulate 4 replicates. In all of our simulations we set the expression level at the root (the most recent common ancestor of all the species) at *β*_*i*_(0) = 1000. We vary *σ*^2^, the variance of the random change due to the Brownian Motion, from 0.1% to 10% of the root expression and use the absolute values of 1, 2, 5, 10, 20, 50, 100. The ratio *γ* of within species variance to *σ*^2^ is varied as 1, 1*/*2, 1*/*4, 1*/*8, 1*/*16, 1*/*32, 1*/*64. We simulated all three modes of gene expression evolution. In the neutral mode of evolution, the rate of attraction to the optimal value in the OU process,*α*, is set to 0 making the gene expression evolve as Brownian Motion only. In the constrained mode of evolution *α* of OU process is positive and all nodes in the phylogenetic tree have the same optimum expression level. In our simulations *α* is varied as 1*/*8, 1*/*4, 1*/*2, 1, 2, 4, 8, 16, 32. Finally, in the adaptive mode of evolution, we considered the two-regime scenario where the two sub-genera *Sophophora* and *Drosophila* are assumed to converge towards two different optimal expression levels.

To ensure an optimal performance of OUCH (A. A. King and Butler 2009) in our evaluation, we let OUCH use the mean expression of simulated replicates and we refer to this model as OUCH.AV. Both methods, EvoGeneX and OUCH.AV, are evaluated as classifiers distinguishing i) genes with constrained expression evolution from the genes with neutrally evolving expression, and ii) genes with adaptive expression evolution from the genes with non-adaptive expression evolution. In both cases we use a variant of the area under precision recall curve metric which we call auPRC_*R*_ and is defined as the area under the part of auPRC curve for the recall value at most *R*, normalised so that maximum auPRC_*R*_ value is 1. This allows us to summarise how the precision changes as recall increases. Note that auPRC_1_ is the standard *auPRC*. We note that OUCH software can also be used to test for simultaneous selection of multiple traits by modeling those traits as a multivariate trait. We asked if this mode can be used to account for replicates by modelling their expressions as a multivariate trait. However, we found that such approach produced results hardly better than a random method in all of our test settings (see Supplementary Section S4 and Supplementary Fig. S7).

### Constrained evolution

Figure 2a shows the auPRC_*R*_ of the two methods under the cutoff values R=0.01, 0.05, 0.1, 0.25, 0.5, 1 for all tested values of *α*. The plots for each *α* aggregate the simulations for all *σ*^2^ and *γ* such that *σ*^2^*/*2*α* ≤ *γσ*^2^, as the within species variance *γσ*^2^ is assumed not to exceed asymptotic between-species variance *σ*^2^*/*2*α* (Bedford and Hartl 2009). Overall, EvoGeneX outperforms OUCH.AV, though the improvement reduces as *α* increases. The higher the alpha, the expression becomes more constrained that it becomes easier for the two models to differentiate from the neutrally evolved expression values.

**Figure 2:**
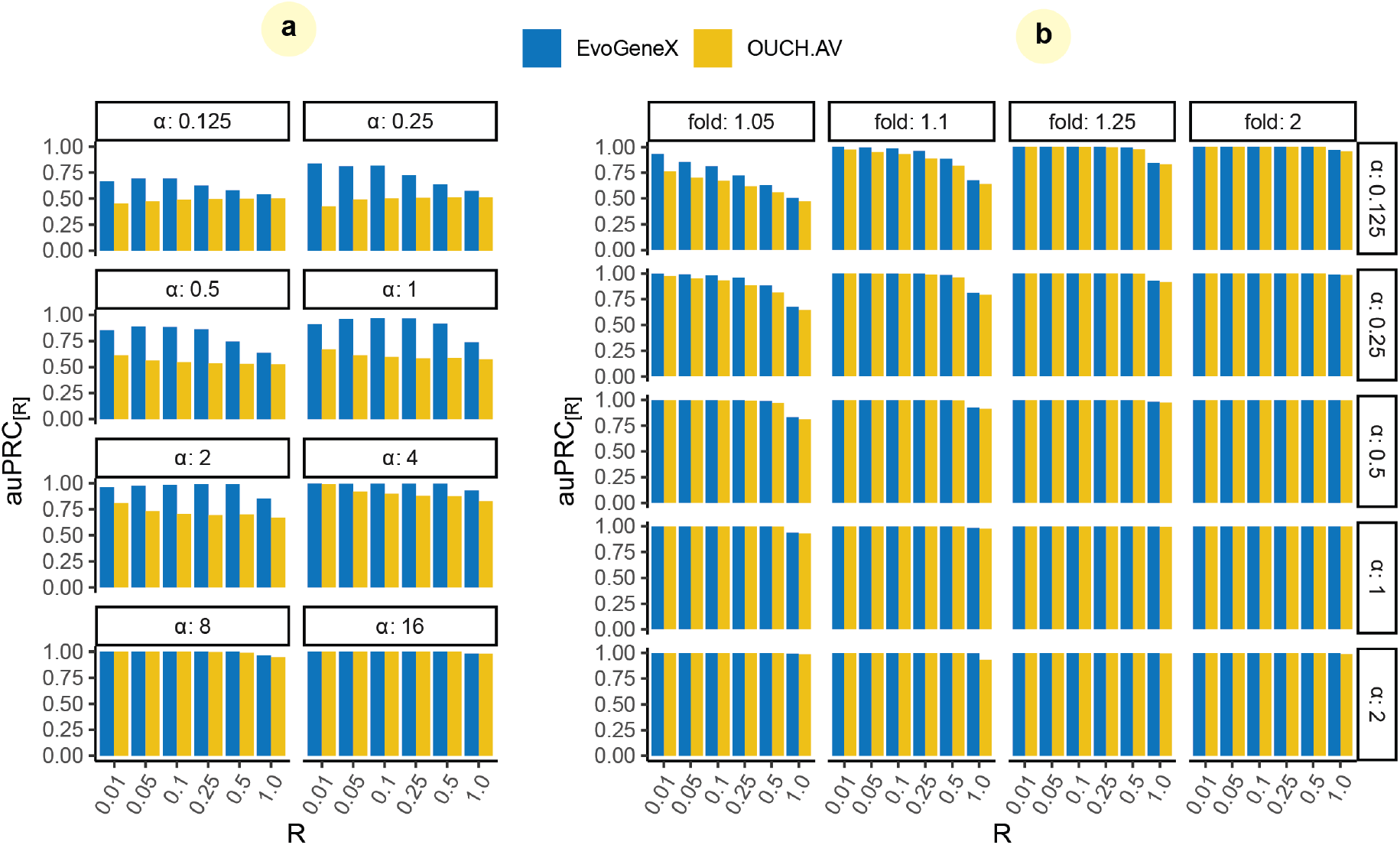
Performance of the methods on simulated data. Shown are values area under Precision Recall Curve (auPRC) at recall cut-off *R*. That is, auPRC*R* is the area under the auPRC curve for the recall value at most *R*, normalised so that maximum auPRC*_R_* value is 1. Each panel corresponds to different parameters as of the simulation given within the white boxes. (**a**) Results on constrained evolution simulations. (**b**) Results on adaptive evolution simulations.

### Adaptive evolution

Previous studies indicate that OUCH-like models can detect adaptive evolution if the separation between optimal values for each regime is large (Cressler et al. 2015). Thus for the simulation of adaptive evolution, we also vary an extra parameter ‘fold’, the ratio of the optimum OU level in the *Drosophila* subgenus to that in *Sophophora* subgenus, as 0.5, 0.8, 0.9, 0.95, 1.05, 1.1, 1.25, 2. The scenarios for the two fold values 0.5 and 2 are symmetric to each other, so for 0.8, 1.25, and so on. Figure 2b shows the *auPRC*_*R*_ of the two methods for the fold values greater than 1. Both the methods are able to correctly detect the adaptive genes when the separation between two optima is larger due to relatively big fold or *α*, however, EvoGeneX works better for the harder cases when both fold and *α* are low. Supplementary Figure S8 shows the same plots with similar trend for all fold values that we experimented with.

Overall, EvoGeneX achieves better auPRC in detecting constrained and adaptive genes especially in the more difficult cases when selection/adaptation is not very strong. In addition, despite the need of estimating an additional parameter and considering larger input data, with more complex relations between them, EvoGeneX is only about 2-fold slower than OUCH to infer from 4 replicates (Supplementary Section S8 and Supplementary Fig. S16).

### 2.3 Stabilising selection and neutral gene expression evolution in *Drosophila*

In this study, we used expression data (Yang et al. 2018) from 9 *Drosophila* species (Fig. 1a): *D. melanogaster* (Dmel), *D. yakuba* (*Dyak*), *D. ananassae* (*Dana*), *D. pseudoobscura* (*Dpse*), *D. persimilis* (*Dper*), *D. willistoni* (*Dwil*), *D. virilis* (*Dvir*), *D. mojavensis* (*Dmoj*) and *D. grimshawi* (*Dgri*) across both sexes and dissected into five body parts: head, gonads, thorax, viscera, and abdomen (see Supplementary Fig. S2 for an illustrative cartoon of the tissues) and a set of 8591 genes with one-to-one orthology in each species as determined in Yang et al. (2018) using multiple evidences.

Given the large evolutionary distances within the *Drosophila* genus, we first asked if the main observations about tissue-specific gene expression in mammals, where the distances are smaller, also hold for *Drosophila*. In particular, it has been observed that when the expression data across different tissues are clustered together the samples cluster first by tissue (Brawand et al. 2011; Chan et al. 2009; J. Chen et al. 2019; Sudmant et al. 2015). We found that this observation also holds for Drosophila (Fig. 3a) with one interesting caveat. Specifically, when we used log gene expression values and Euclidean distances then head and thorax of the same specie often clustered together (Supplementary Figure S6). This co-clustering may reflect the fact that in fly these two anatomical regions have extensive central nervous system components (brain and thorasic ganglia). Thus both tissue-specific and species-specific trends can be observed, depending on the weight given to genes in the tails of the expression distributions. In addition, studies in mammals (J. Chen et al. 2019) report that expression evolution within individual tissues generally follows species evolution. We confirmed that this is also the case for *Drosophila* (Fig. 3c and Supplementary Table S2). In particular, we could reconstruct the phylogenetic tree (either perfectly or with small Robinson-Foulds distances from the known sequence based tree) using gene expression values in any specific body part in either sex. Thus, while the evolution of gene expression is consistent with species evolution, the fact that all samples cluster predominately by body parts and in some cases by sex, suggests that gene expression evolution is also subject to strong tissue and sex specific constraints.

**Figure 3:**
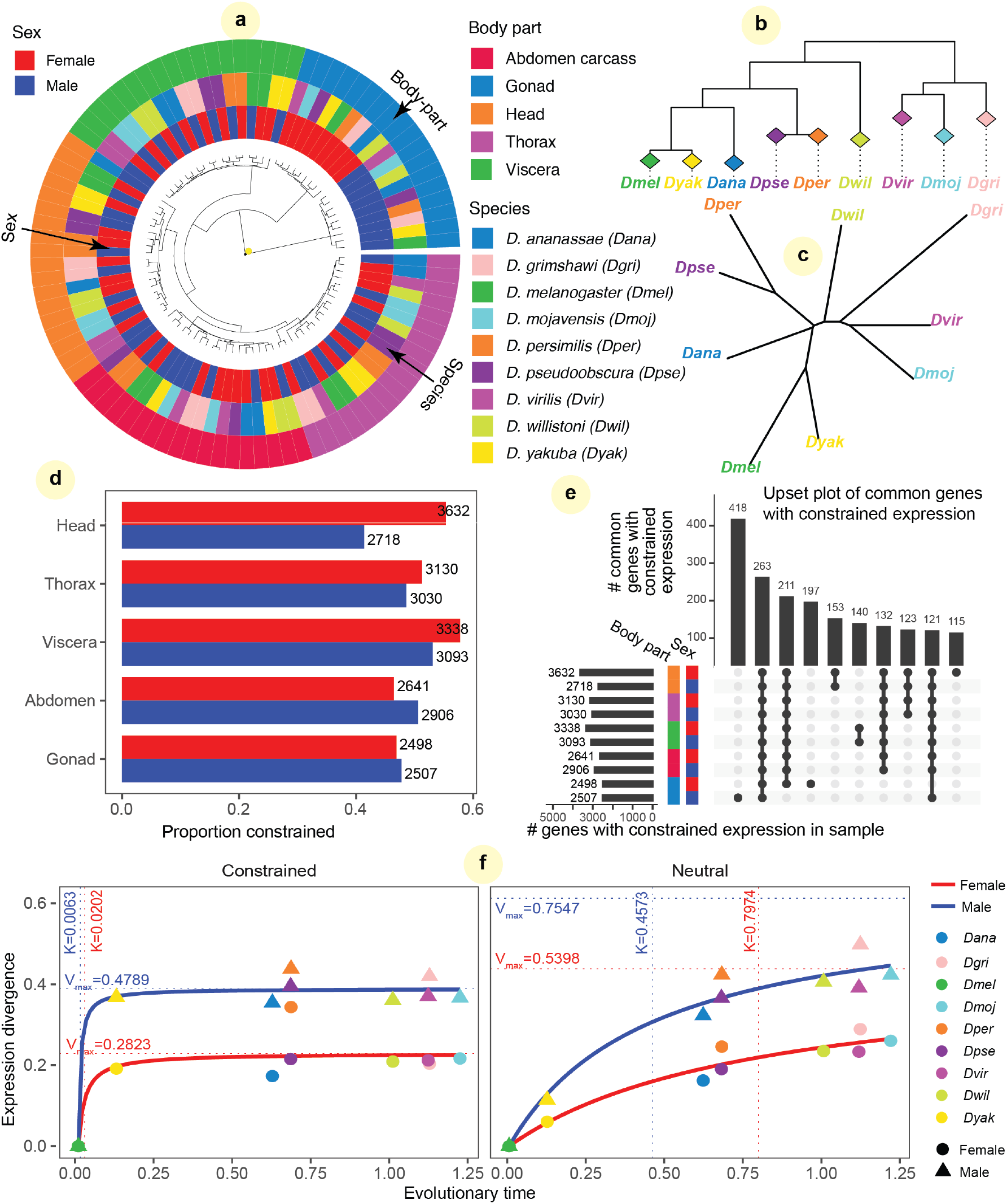
Constrained and neutral gene expression evolution in *Drosophila* **genus**. (**a**) Hierarchical clustering of gene expression data (using 1-Spearman’s correlation). The outside ring represents body parts, the middle ring species, and the inside ring sex. The main organizing factor is body part (except for gonads which cluster first by sex). (**b**,**c**) Neighbour Joining trees constructed using gene expression similarity in specific body parts (the tree shown is for female head) illustrate that gene expression evolution (**c**) typically follows the species evolution (**b**). (**d**) The proportions of the number of genes with constrained expression evolution to the number of genes for which neutral expression evolution could not be rejected. (**e**) UpSet plot (Lex et al. 2014) showing groups of common genes that have constrained expression in a subset of body parts and sexes (black circles connected by vertical line). The groups are ordered by the decreasing number of common genes and 10 most abundant groups are shown. (**f**) Modeling evolutionary dynamics in female gonads (ovaries) with the Michaelis-Menten curve. *Vmax* estimates equilibrium divergence and *K* denotes the time to reach half of *Vmax*. Constrained genes are characterized by smaller *K* and smaller *Vmax*.

Next, we used EvoGeneX to identify genes that are subject to constrained expression evolution and those for which the null hypothesis of neutral expression evolution could not be rejected. To focus on the genes that are relevant for a given tissue and sex, we included in this analysis only those genes that have normalised read counts of more than one in all 4 replicates and all species. Independent of organ and sex, and despite the stringency of our model, the hypothesis of neutral evolution could be rejected for more than 40% of the expressed genes (Fig. 3d). In the following we refer to a gene as “constrained” if the null hypothesis was rejected for that gene, and “neutral” otherwise. Note that, technically, this neutral group of genes with drifting expression also contains genes with weakly constrained expression which could not be confidently distinguished from the neutrally evolving group (analogous to nearly neutral sequence evolution (Ohta 1973)). Interestingly we found that gene expression evolution in females was often more constrained than in males with biggest differences in head and viscera. As discussed in the previous section, gene expression evolution for individual body parts and sexes follows the previously estimated sequence based tree species tree (Z.-X. Chen et al. 2014) either perfectly or with small Robinson-Foulds distances. We asked if the occasional small differences between the sequence based tree and expression based trees would be eliminated if we use only the genes with neutral (or nearly neutral) expression evolution in the give body part and sex only. We found this indeed be the case although some differences persisted (Supplementary Table S2).

It is widely accepted that genes performing essential cell functions show constrained sequence evolution (Fisher 1930). Therefore, we asked if genes with constrained expression in one body part or sex show constrained expression in other body parts or in the opposite sex. Interestingly, the largest set of genes with constrained expression in more than one body part occurred in the intersection of all the body parts independent of the sex (Fig. 3e for overlaps of more than 100 genes and Supplementary Fig. S9 for more overlaps).

Gene Ontology Enrichment Analysis (GOEA, Table 1) revealed that the most significantly enriched GO term for biological process for this common gene-set is mRNA splicing, via spliceosome. Alternative splicing is a highly conserved mechanism common to eukaryotes acting to modify expression of specific isoforms in specific spatio-temporal patterns. We also considered gene expression constrained in smaller subsets of body parts and sexes. For example, genes with constrained expression only in the head of both male and female were enriched in the GO terms for visual perception. Vision is an important evolutionary trait, and thus, many associated genes are indeed expected to undergo purifying selection. Finally, consistent with the fact that fly head and thorax share extensive central nervous system components, the common genes with constrained expression exclusively in those two body parts were found to have enriched GO terms for nervous system process and dendrite development.

**Table 1:**
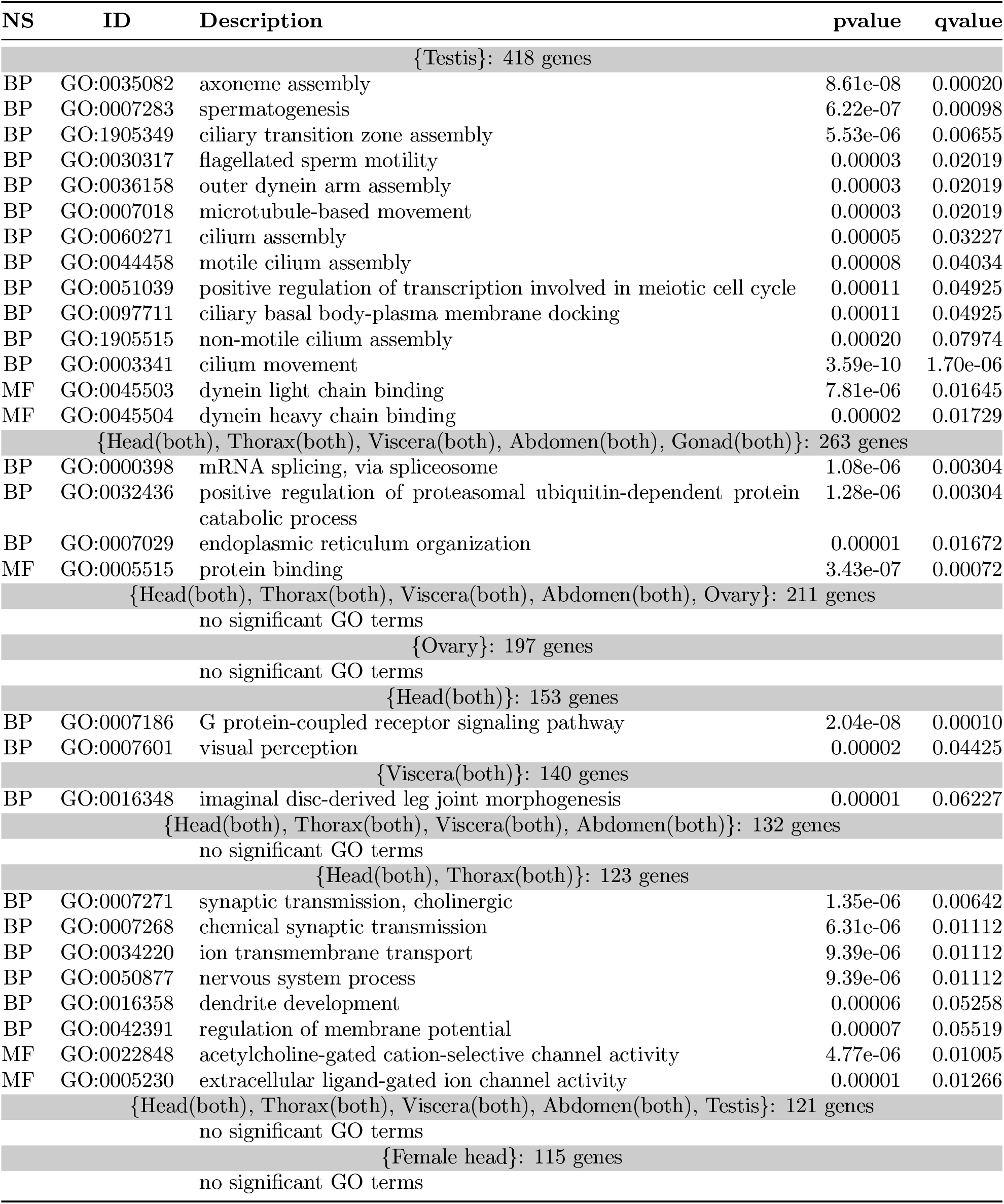
Gene Ontology Enrichment Analysis of the common constrained genes

| NS | ID | Description | pvalue | qvalue |
| --- | --- | --- | --- | --- |
| {Testis}: 418 genes |  |  |  |  |
| BP | GO:0035082 | axoneme assembly | 8.61e-08 | 0.00020 |
| BP | GO:0007283 | spermatogenesis | 6.22e-07 | 0.00098 |
| BP | GO:1905349 | ciliary transition zone assembly | 5.53e-06 | 0.00655 |
| BP | GO:0030317 | flagellated sperm motility | 0.00003 | 0.02019 |
| BP | GO:0036158 | outer dynein arm assembly | 0.00003 | 0.02019 |
| BP | GO:0007018 | microtubule-based movement | 0.00003 | 0.02019 |
| BP | GO:0060271 | cilium assembly | 0.00005 | 0.03227 |
| BP | GO:0044458 | motile cilium assembly | 0.00008 | 0.04034 |
| BP | GO:0051039 | positive regulation of transcription involved in meiotic cell cycle | 0.00011 | 0.04925 |
| BP | GO:0097711 | ciliary basal body-plasma membrane docking | 0.00011 | 0.04925 |
| BP | GO:1905515 | non-motile cilium assembly | 0.00020 | 0.07974 |
| BP | GO:0003341 | cilium movement | 3.59e-10 | 1.70e-06 |
| MF | GO:0045503 | dynein light chain binding | 7.81e-06 | 0.01645 |
| MF | GO:0045504 | dynein heavy chain binding | 0.00002 | 0.01729 |
| {Head(both), Thorax(both), Viscera(both), Abdomen(both), Gonad(both)}: 263 genes |  |  |  |  |
| BP | GO:0000398 | mRNA splicing, via spliceosome | 1.08e-06 | 0.00304 |
| BP | GO:0032436 | positive regulation of proteasomal ubiquitin-dependent protein catabolic process | 1.28e-06 | 0.00304 |
| BP | GO:0007029 | endoplasmic reticulum organization | 0.00001 | 0.01672 |
| MF | GO:0005515 | protein binding | 3.43e-07 | 0.00072 |
| {Head(both), Thorax(both), Viscera(both), Abdomen(both), Ovary}: 211 genes |  |  |  |  |
| no significant GO terms |  |  |  |  |
| {Ovary}: 197 genes |  |  |  |  |
| no significant GO terms |  |  |  |  |
| {Head(both)}: 153 genes |  |  |  |  |
| BP | GO:0007186 | G protein-coupled receptor signaling pathway | 2.04e-08 | 0.00010 |
| BP | GO:0007601 | visual perception | 0.00002 | 0.04425 |
| {Viscera(both)}: 140 genes |  |  |  |  |
| BP | GO:0016348 | imaginal disc-derived leg joint morphogenesis | 0.00001 | 0.06227 |
| {Head(both), Thorax(both), Viscera(both), Abdomen(both)}: 132 genes |  |  |  |  |
| no significant GO terms |  |  |  |  |
| {Head(both), Thorax(both)}: 123 genes |  |  |  |  |
| BP | GO:0007271 | synaptic transmission, cholinergic | 1.35e-06 | 0.00642 |
| BP | GO:0007268 | chemical synaptic transmission | 6.31e-06 | 0.01112 |
| BP | GO:0034220 | ion transmembrane transport | 9.39e-06 | 0.01112 |
| BP | GO:0050877 | nervous system process | 9.39e-06 | 0.01112 |
| BP | GO:0016358 | dendrite development | 0.00006 | 0.05258 |
| BP | GO:0042391 | regulation of membrane potential | 0.00007 | 0.05519 |
| MF | GO:0022848 | acetylcholine-gated cation-selective channel activity | 4.77e-06 | 0.01005 |
| MF | GO:0005230 | extracellular ligand-gated ion channel activity | 0.00001 | 0.01266 |
| {Head(both), Thorax(both), Viscera(both), Abdomen(both), Testis}: 121 genes |  |  |  |  |
| no significant GO terms |  |  |  |  |
| {Female head}: 115 genes |  |  |  |  |
| no significant GO terms |  |  |  |  |

### 2.4 Differences in evolutionary dynamics across sexes and organs

Next, we wanted to compare the evolutionary dynamics of gene expression between the sexes and the body parts. To do this, we considered the relation between the evolutionary distance (as determined by sequence evolution) of a species with respect to a reference, say *D. melanogaster*, and the expression divergence measured as 1 *− S*(*dmel, X*) where *S*(*dmel, X*) is some measure of similarity, say the square of Spearman’s correlation, between the replicate-averaged expressions in *D. melanogaster* and species *X* for a group of genes. Among closely related taxa such expression divergence is expected to be a linear function of time, but on larger distances it becomes nonlinear and saturates on a level that depends on the selection strength (Bedford and Hartl 2009; Whitehead and Crawford 2006) and potentially other biological limitations. This type of kinetics is similar to classical enzymatic kinetics, and can be well approximated by the Michaelis-Menten curve *f* (*x*) = *xV*_*max*_*/*(*K* +*x*). Figure 3f shows such curves for the constrained and neutrally evolving gene sets separately in the gonads of either sexes. The parameter *V*_*max*_ is the maximum (putative equilibrium value) of 1 *− S*(*dmel, X*) and the parameter *K* is the value of *x* where *f* (*x*) = *V*_*max*_*/*2. Thus *V*_*max*_ estimates the constraints on the divergence of the given group of genes while *K* measures the ‘speed’ of evolutionary dynamics: *K* close to zero corresponds to the situation where the value *V*_*max*_ corresponding to the equilibrium divergence is immediately achieved. In contrast, larger *K* corresponds to slower relative divergence to the putative equilibrium at *V*_*max*_. The values *V*_*max*_ and *K* can be used to compare the evolutionary dynamics of gene expression between different organ systems, or gene groups.

Table 2 shows the values of *V*_*max*_ and *K* for the two gene groups for all combinations of tissues and sexes (Supplementary Fig. S10 shows the fitted curves). As expected, *V*_*max*_ values for the groups of genes with constrained expression evolution are systematically smaller than the *V*_*max*_ values of gene with neutrally evolving expression for the same tissue/sex, although in the case of abdomen carcass the difference is small (consistent with the small *K* values for neutral expression evolution in abdomen carcass). Strikingly, male gonads show far less constrained expression evolution (larger *V*_*max*_) than any other body part in any sex. This is true not only for the constrained expression group, but also the high *V*_*max*_ for genes with neutrally evolving expression in male gonads. The testes show the highest sex-biased expression in Drosophila (Parisi et al. 2004; Arbeitman et al. 2002) and it has been observed that the evolution of genes with sex-biased expression is accelerated (Haerty et al. 2007; Begun et al. 2007; Brown et al. 2014; Z.-X. Chen et al. 2014; Zhang et al. 2007; Meisel 2011).

**Table 2:**
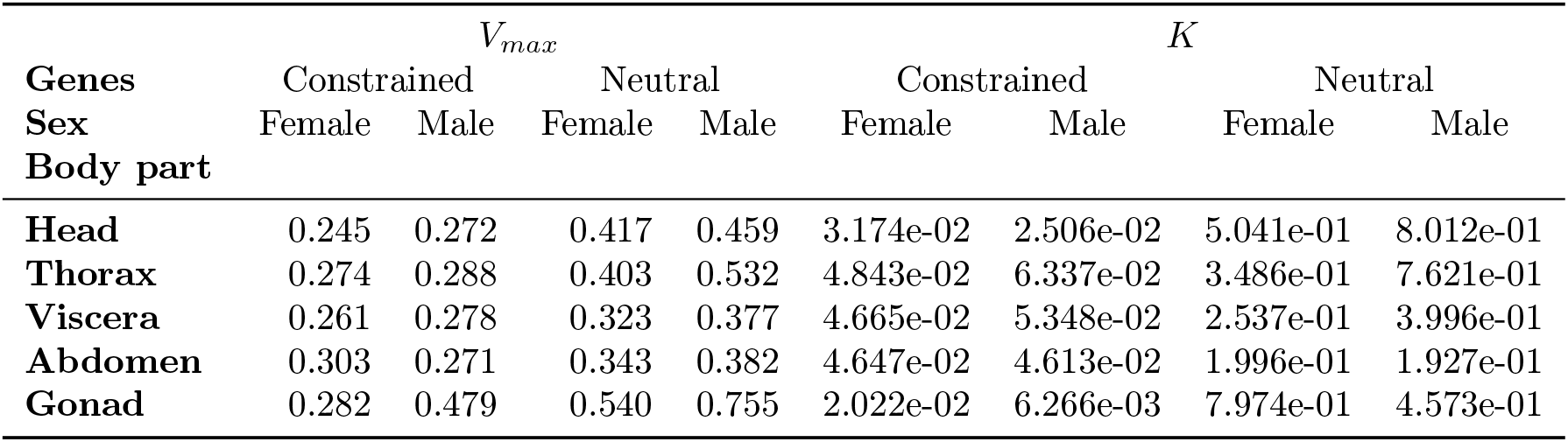
Summary of *V*_*max*_ and *K* of constrained and neutral genes for different tissues and sexes

The high *V*_*max*_ value for neutrally evolving genes in female gonad is surprising since, unlike male, female gonad gene expression is not particularly less constrained than other organ systems. Indeed, the large *K* value for neutrally evolving female gonad gene expression suggests that the changes in gene expression (relative to the *V*_*max*_ value) are slow. These results show that the expression evolution of male and female gonads follow drastically different dynamics. Importantly, these results do not depend on the species taken as reference (*D. melanogaster* here) as we observe similar results when *D. pseudoobscura* or *D. virilis* is taken as reference (Supplementary Fig. S11 and Table S3). These appear to be general features of gene expression evolution in the phylogeny.

### 2.5 EvoGeneX reveals examples of adaptive evolution

Differences in habitat, reproductive strategy, and other factors should lead to evolutionary shifts of optimal expression values for some genes in a particular sex and/or organ system. As a result, the optimum value of a gene expression for a given sex and organ system might be different in different branches of the evolutionary tree.

Statistically, detection of such evolutionary adaptation is very challenging and current methods have limited statistical power that has been attributed to the relatively small sizes of evolutionary trees for which tissue-specific gene expression data is available (J. Chen et al. 2019). We reasoned that by utilizing replicates for the estimation of within-species variability, EvoGeneX might have greater power to detect such adaptive evolution of gene expression. We used a hypothesis testing framework, and tested if both the neutral BM model and constrained model (OU with a single optimum expression) are rejected in favor of adaptive evolution model (OU with multiple optima). We tested for evidence of adaptive evolution in two main settings. First we use a two-regime setting where the regimes correspond to the two sub-genera: *Sophophora* and *Drosophila* (Fig. 1a). In addition, to capture potential adaptation in smaller subgroups of species, we also consider a three-regime setting where we partition the tree into three regimes of equal number of species (*D. melanogaster, D. yakuba, D. ananassae*), (*D. pseudoobscura, D. persimilis, D. willistoni*), (*D. virilis, D. mojavensis, D. grimshawi*). We call the first and the third group as *Melanogaster* and *Drosophila*, respectively, and the remaining species containing *Obscura* group and *D. willistoni* as *Obscurawil* due to the lack of standard nomenclature for such a group (Supplementary Fig. S1).

Interestingly, in the two-regime scenario, the highest number of genes with adaptive expression evolution was observed in male head (Fig. 4a). For the three regime scenario, in addition to male head, high numbers of genes with adaptive expression evolution were observed in female gonads (Fig. 4a). Expression adaptation in male heads might be related to the fact that males have extensive sets of species-specific mating behaviors ranging from singing to displaying in mating arenas (A. Singh and B. N. Singh 2014). We also note that there are differences in wiring and gross anatomy between male and female fly brains (Cachero et al. 2010). Drosophila ovaries also vary in morphology (Mahowald and Kambysellis 1980). The set of adaptive genes uncovered by EvoGeneX can provide additional cues for sex specific adaptation and warrants further investigation.

**Figure 4:**
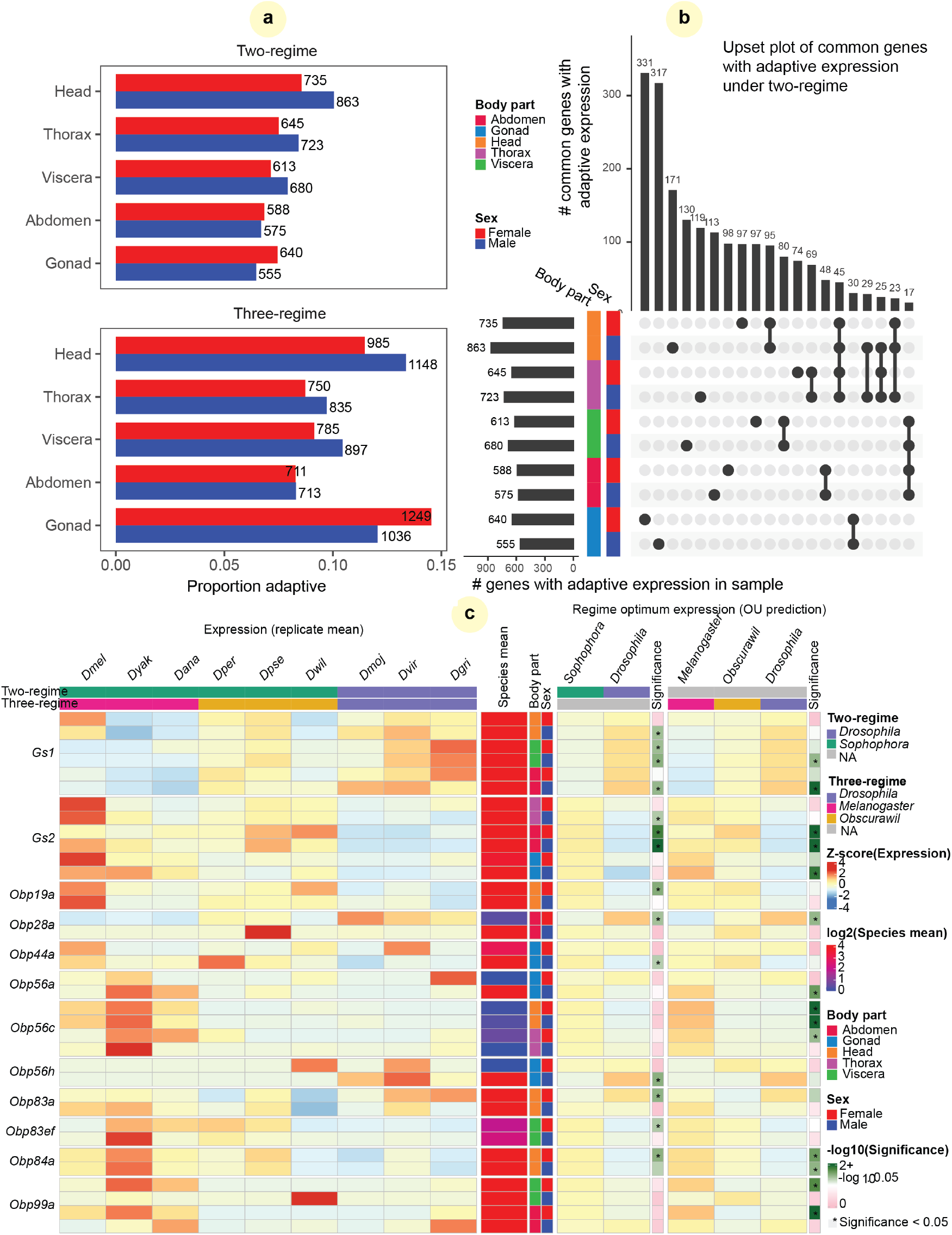
Adaptive gene expression evolution in *Drosophila* **genus**. (**a**) The proportion and number of adaptive genes in each body parts and sex for the two partition into evolutionary regimes considered in the study. (**b**) UpSet plot showing number of common genes with adaptive expression evolution in two-regime (see Fig. 3e for full description). (**c**) Examples of adaptive genes. Shown from left to right are: z-score of gene expression in the given body part and sex, average expression for the region and sex, color code for body part and sex (here we show both sexes for any body part where adaption was detected, see Supplementary Fig. S15 for all body parts), the optimal expression values obtained for each regime by EvoGeneX in the two-regime scenario and the max p-values for rejecting neutral and constrained evolution. Next, the same information is shown for the tree-regime scenario.

Interestingly, in contrast to the constrained expression expression evolution, where many genes with constrained expression were common to all the body parts (Fig. 3e), the adaptive evolution is largely body part specific.

It has been suggested that one important mechanism facilitating adaptation is gene duplication. Gene duplication is usually followed by degeneration of one copy, but sometimes each of the duplicates evolves a specialized function. There are two Glutamine Synthase encoding genes in the Drosophila genus (encoded by *Gs1* and *Gs2*). These genes play an important role in nitrogen metabolism, amino acid synthesis, neurotransmiter recycling, and amonia detoxification in the mitochondria (Gs1) and cytoplasm (Gs2) (Caizzi et al. 1990; Vernizzi et al. 2020). We found that *Gs1* expression was elevated, while reciprocally *Gs2* was depressed in the Drosophila subgenus, raising the possibility of compensatory expression evolution of these enzyme encoding genes (Fig. 4c). One important element of adaptation is sensing the environment, including food sources, and mates. Therefore, we focused on the expression of odor binding proteins (Obp) as potential examples of adaptive change. *Obp* genes are members of a gene family evolved by duplication that has extensively diversified. While the exact molecular role of a large number of Obps is not completely clear, they do play a role in pheromone-dependent species and sex recognition (Xiao et al. 2019). For example, *D. grimshawi* males paint attractive pheromones onto the substrate in mating arenas, known as leks, to attract competing males and female observers. Additionally both *D. virilis* and *D. mojavensis* use aggregation pheromones to mark resources for cooperative exploitation. Expression could come under selection for higher expression in opposite sexes due to a switch in the role of a particular odor and/or hormone from intra-sexual to extra-sexual, or between females and males. Indeed, we found that expression of many genes encoding odor binding proteins are subject to adaptive expression evolution; often in a sex-biased manner (Fig. 4c). As expected, such adaptive expression evolution of odor binding proteins is mostly found in head (where olfactory organs are located) but sometimes in other body parts as well (Fig. 4c). For example, the Obp28a protein binds floral odors (Gonzalez et al. 2020) and we found higher adaptive expression of *Obp28a* in Drosophila subgenus female abdomens. Additionally, while we haven’t detected statistically significant adaptation in males within the tested regimes, we saw a species-specific increase in *D. pseudoobscura* male abdomen. Next, we observed high adaptive expression of *Obp56a* in the melanogaster group testis. Expression of *Obp56a* is known to be downregulated in mated female *D. melanogaster* (McGraw et al. 2004). *Obp56c* expression showed adaptive increases in melanogaster group heads in the three regime analysis. Knockdown of *Obp56h* results in rapid male remating and those males produce less 5-tricosene, an anti-aphrodisiac that inhibits male-male courtship (Shorter et al. 2016). Interestingly, we found *Obp56h* to be adaptive in testis. Very little is known about the *Obp83* and *Obp84* gene families or *Obp19a*, but these showed striking female-specific adaptation.

## 3 Conclusions

Natural selection is a complex organism level force that has well studied effects on the sequence of genes, gene duplication and divergence, and *de novo* gene formation. But, because all cell types have the same genome, the genome sequence is always a compromise. Sequence is selected based on the survival of a genotype, while genes are almost always expressed in multiple contexts within an organism and are thus free to optimize expression levels in different cell types. Thus studies of the evolution of gene expression tuning can contribute to understanding the molecular basis of phenotypic traits.

Early in the genomic era, biological replication was not often a priority. The steadily decreasing cost of sequencing now motivates researchers to measure gene expression in multiple biological replicates. Our own data utilised in this study, includes 4 replicates per each specie, sex and body part. This data allowed us to estimate within species expression variability. We started with the assumption that information about within-species variation should boost the performance of the OU-based methods without increasing the number of species under study. We then developed a new method, EvoGeneX which formally includes within-species variation estimated using biological replicates. Overall, our results demonstrate that, by taking full advantage of the existing gene expression data, EvoGeneX provides a new and more powerful tool to study gene expression evolution relative to the currently leading method.

There have been several studies of Drosophila expression evolution as it relates to sex, protein coding sequence, and expression breadth (Meisel 2011; Ellegren and Parsch 2007; Campos et al. 2018). Some of those studies have been contradictory or focused on a single prevalent pattern of divergence rather than distinct evolutionary mechanisms. We used EvoGeneX to provide a mathematical framework for an extensive analysis of sex and body part specific gene expression evolution in the *Drosophila* genus. In agreement with some of the previous studies, we showed that neutral gene expression evolution can be confidently rejected in favor for constrained evolution (presumably due to purifying selection) in nearly half of the genes. However it also showed that for a comparable number of genes the neutral expression evolution could not be rejected indicating either truly neutral expression evolution or expression evolution under very weak constraints. In addition, our analysis uncovered interesting but differential adaptive evolutionary dynamics especially for male head and ovaries. Overall, testis showed weaker constraints on gene expression evolution (although the number of constrained genes was comparable between male and female) but at the same time were overall less adaptive. In contrast, in male head we discovered more genes showing adaptive gene expression evolution relative to female head.

Using EvoGeneX, we also detected interesting examples of adaptive evolution including prominent examples of odor biding proteins as well as an example of adaptive evolution followed gene duplication. Interestingly, in contrast to the constrained expression expression evolution where many genes with constrained expression were common to all the body parts, we found that the adaptive evolution tends to be body part specific.

In summary, our tool performs well to uncover data showing that gene expression is an important trait under selection and subject to drift at the body part level, to optimize the survival and reproduction of the individual – the unit of selection. This will likely also be true at the cell type level, and it is likely that data allowing testing this hypotheses will soon be available.

## 4 Materials and Methods

### Data and data / software availability

The expression data (Yang et al. 2018) was obtained from the NCBI GEO database under accession numbers GSE99574 and GSE80124, and the phylogenetic tree was obtained from Z.-X. Chen et al. (2014) (Supplementary Section S1). The source code for EvoGeneX is available at the NCBI public Github repository: https://github.com/ncbi/EvoGeneX. A software pipeline to analyze the data using EvoGeneX was built using JUDI (Pal and Przytycka 2019).

### EvoGeneX model and its parameters

EvoGeneX takes as its input a rooted evolutionary tree and the values of quantitative characters *y*_*i,k*_ for all terminal taxa *i* and biological replicates *k*. Two sets of random variables, *X*_*i*_(*t*) at the taxa level and *Y*_*i,k*_ at the replicate level, are used to model the evolution of trait value across time such that the observed trait value at time *T*_*i*_, the evolutionary time from the least common ancestor of all species in this tree to species *i*, is *Y*_*i,k*_(*T*_*i*_) = *y*_*i,k*_. The two sets of random variables are governed by equations (2) and (3).

EvoGeneX further assumes that the optimum value *β*_*i*_(*t*) of “attraction” in (3) changes at speciation events only and remains constant along individual edges of the phylogenetic tree. The history of the *i*th lineage consists of a number, *κ*(*i*), of sequential branch segments demarcated by speciation events 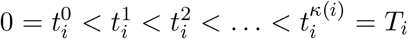. Let all 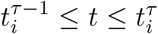 represent a *selective regime* where the evolution “attracts” towards a fixed optimum value 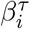 of *β*_*i*_(*t*). EvoGeneX further simplifies the model by letting a small number, *R*, of distinct optimum values *θ*_*r*_, *r* = 1, …, *R*, each corresponding to one selective regime. In fact one of the most interesting cases corresponds to the model with two optima where one branch of the tree follows a regime of optimum values *θ*_1_ and the rest of the tree *θ*_0_ (J. Chen et al. 2019; Brawand et al. 2011).

Let the binary variable 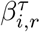 indicate if the *τ* th branch on lineage *i* has operated in *r*th regime. Then we have 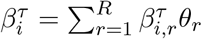. Since each branch is associated with exactly one optimum, for each *i, τ* there is exactly one *r* such that 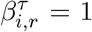 and 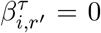 for all *r* ≠ *r* ′. Further, self-consistency requires that 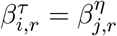 whenever lineage *i* and *j* share the branch ending in epoch 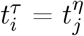.

Thus, for a given tree, *y*_*i,k*_s and 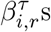, EvoGeneX estimates the parameters *α, σ, γ, θ*_0_, *θ*_1_, …, *θ*_*R*_.

### Inference of EvoGeneX parameters using Maximum Likelihood estimates

In the following, it will be convenient to make use of matrix notation. Accordingly, we collect our random variables, *X*_*i*_(*t*) for the trait values at the taxa, and *Y*_*i,k*_(*t*) for the replicated trait values, in vectors **x**(*t*) and **y**(*t*), respectively, and our observed quantitative data in vector **y** with components *y*_*i*+(*k−*1)*N*_ = *Y*_*i,k*_(*T*_*i*_), the observed value of replicate *k* of taxa *i* at the evolutionary time *T*_*i*_. The expected value of random variable **y**(*t*) at the taxa can be shown to be (Supplementary Section S3)

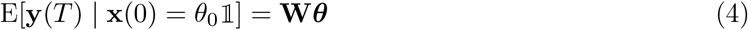

where 𝟙 is a vector of all 1s, column vector ***θ*** = (*θ*_0_, *θ*_1_, …, *θ*_*R*_)^*T*^ and the weight matrix *W* is dependent only on *α* among the parameters and has entries

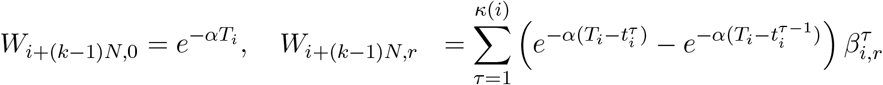

for *i* = 1, …, *N, k* = 1, …, *M* and *r* = 1, …, *R*. Similarly, let an *MN × MN* matrix **V** give the covariance between species *i*, replicate *k* and species *j*, replicate *l* by the entry (Supplementary Section S3)

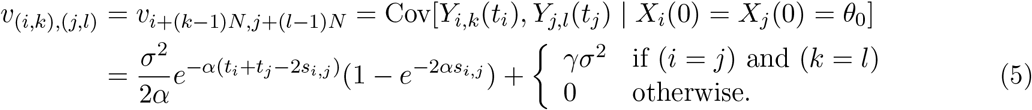

It is known that **y** is a multi-variate Gaussian ∼ 𝒩 (**W*θ*, V**) with mean and co-variance given by equations (4) and (5) (Hansen and Martins 1996). Thus, the likelihood of the parameters *α, σ, γ*, and ***θ***, given the data **y** is

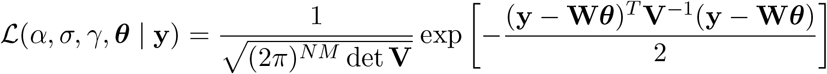

As maximizing *ℒ* is equivalent to minimizing *U* = *−*2 log *ℒ*, we seek to minimize

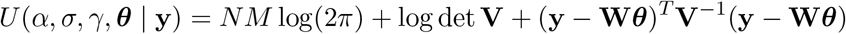

However, it can be noted that **V** has a nice structure and can be expressed as 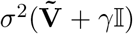 where 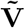 is dependent on *α* only among all the parameters and 𝕀 is an identity matrix of size *MN* .The elements of 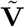 are given by 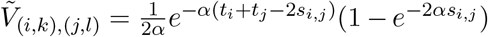. Thus, *U* can be expressed as

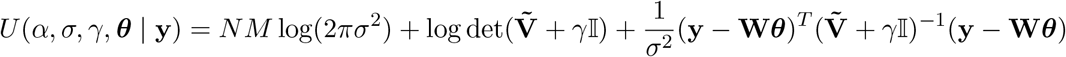

whose minimum can be estimated using any off-the-self nonlinear optimization solver.

However, we improve efficiency by utilizing Karush-Kuhn-Tucker conditions. By setting the partial derivatives of *U* with respect to *σ* and ***θ*** to 0 at an optimal solution 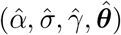, we get

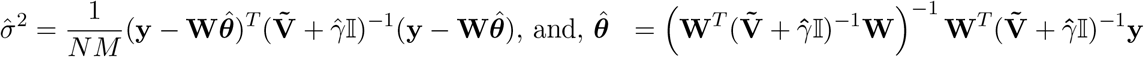

Thus, instead of minimizing function *U* of four parameters *α,σ,γ* and ***θ***, it is enough to minimize a new function of two parameters, *α* and 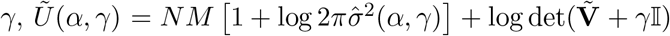 where the following two intermediate functions

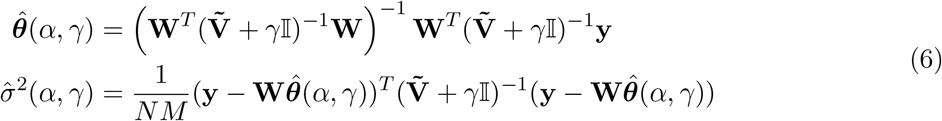

give the values of the remaining two parameters *σ* and ***θ*** at the optimal solution.

### Maximum Likelihood estimates for Brownian Motion model

In order to compare EvoGeneX model of evolution with the BM model, we need to compute maximum likelihood estimates for the Brownian Motion model (BM) accounting for the within-species variation. BM is a simplified model in comparison to EvoGeneX: there is no “attracting” optimal values and hence there is no *α* parameter and *θ* has only one value, *θ*_0_, to be estimated. Using a procedure similar to the previous section, we estimate *σ, γ, θ*_0_ from the given data (Supplementary Section S3).

### Computing statistical significance

We use statistical hypothesis testing to decide which of the three different modes of evolution the trait has undergone: i) neutral, ii) constrained and iii) adaptive (Section 2.1). For this purpose we use the likelihood ratio test (Supplementary Section S3).

## Acknowledgement

The authors would like to thank Haiwang Yang and Justin Fear, for their helpful comments. This research was supported by the Intramural Research Program of the National Library of Medicine and the National Institute of Diabetes and Digestive and Kidney Diseases at the National Institutes of Health, USA.

