## Supplemental Information for "Modeling gene expression evolution with EvoGeneX uncovers differences in evolution of species, organs and sexes"

#### S1 Data

##### S1.1 Phylogenetic tree

We obtained the phylogenetic tree for 12 *Drosophila* species (Z.-X. Chen et al. 2014) in Newick format via personal communication. The tree was trimmed to 9 *Drosophila* species of our interest using `drop.tip` function of `ape` R package. The trimmed Newick file used in our analysis is reproduced here.

```
((dgri:0.206091,(dmoj:0.235899,dvir:0.134088):0.067592):0.235776,
 (dwil:0.442195,((dper:0.006819,dpse:0.005611):0.262193,
                 (dana:0.316433,(dyak:0.061958,dmel:0.059667):0.244496):0.107103
               ):0.152356
 ):0.117888
);
```

Figure 1a in the main text shows a drawing of the phylogenetic tree. Additionally it shows the branches on which the two levels of 'optimum' expression corresponding to *Drosophila* and *Sophophora* subgenera in the Ornstein-Uhlenbeck model under two-regime are assumed to be active. Figure S1 in this document shows the branch colors according to the 'optimum' levels active on them under three-regime for the subgenra *Melanogaster* and *Drosophila* and the rest of the species that includes *D. willistoni* and the *Obscura* group and we name as *Obscurawil*.

##### S1.2 Gene expression data

The expression data was obtained from (Yang et al. 2018). The data is available at the NCBI GEO database with accession numbers GSE99574 (for non-Hawaiian species) and GSE80124 (for Hawaiian species). Figure S2 shows a cartoon for the nomenclature of different tissues. We took the samples from the body parts for which all 4 replicates were available. These were: head (HD), Thorax (TX) Abdomen Carcass (AC), Viscera (VC) and Gonad (GO). For *D. melanogaster*, we used w1118 strain.

The raw read counts are available in the supplementary folders in the GEO database. The data was normalized to allow for the gene level (between samples) analysis (Yang et al. 2018). Specifically, all body parts and all samples were first normalized using DESeq2 (Love et al. 2014) and then only the gene-level normalized counts for 8591 orthologous genes were used for our evolutionary analysis. The mapping of *D. melanogaster* gene-IDs to the ortholog-IDs in other species were also downloaded from the GEO supplementary folders.

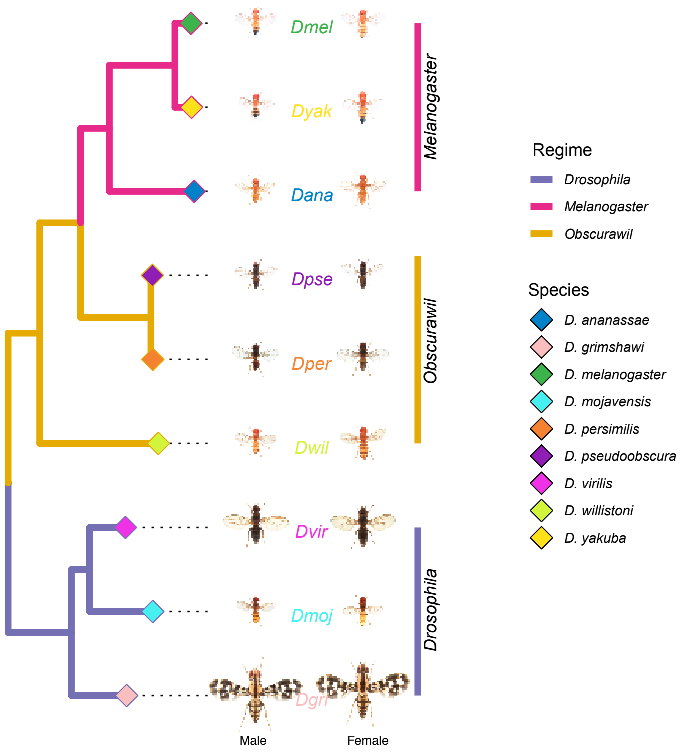


Figure S1: **Three-regime scenario**

### S2 Exploratory analysis of data

#### S2.1 Summary statistics

The expression of the replicates were found to have 'good' correlations. Table S1 shows the Pearson's correlation coefficients for each pair among the 4 replicates for each species, body part and sex. Most of the coefficients are more than 0.9. Figure S3 shows the summary of expression values each replicate and Fig. S4 shows the violin plots.

Table S1: **Pearson's correlation coefficients between any pair of replicate expressions.**

| **Species** | **Body part** | **Sex** | R1vR2 | R1vR3 | R1vR4 | R2vR3 | R2vR4 | R3vR4 |
|---|---|---|---|---|---|---|---|---|
| Dmel | **Head** | **Female** | 0.994284 | 0.990478 | 0.992961 | 0.964538 | 0.972433 | 0.985174 |
| | | **Male** | 0.998499 | 0.996507 | 0.996663 | 0.997282 | 0.996951 | 0.998052 |
| | **Thorax** | **Female** | 0.994681 | 0.993190 | 0.994309 | 0.985519 | 0.990970 | 0.992146 |
| | | **Male** | 0.991482 | 0.993884 | 0.992369 | 0.987842 | 0.992546 | 0.994792 |
| | **Viscera** | **Female** | 0.993888 | 0.997742 | 0.991912 | 0.982094 | 0.993345 | 0.979998 |
| | | **Male** | 0.987480 | 0.984554 | 0.966164 | 0.990095 | 0.987106 | 0.987937 |

Continued on next page

Table S1 – continued from previous page

| Species | Body part | Sex | R1vR2 | R1vR3 | R1vR4 | R2vR3 | R2vR4 | R3vR4 |
|---|---|---|---|---|---|---|---|---|
| | **Abdomen** | **Female** | 0.988949 | 0.984882 | 0.994223 | 0.990497 | 0.993050 | 0.991092 |
| | | **Male** | 0.992132 | 0.980661 | 0.987349 | 0.970172 | 0.979731 | 0.991954 |
| | **Gonad** | **Female** | 0.990568 | 0.979192 | 0.983561 | 0.991353 | 0.995576 | 0.977971 |
| | | **Male** | 0.999377 | 0.999368 | 0.999392 | 0.998940 | 0.999083 | 0.999232 |
| **Dyak** | **Head** | **Female** | 0.984920 | 0.977258 | 0.996401 | 0.983580 | 0.988325 | 0.991220 |
| | | **Male** | 0.995152 | 0.983804 | 0.981935 | 0.993153 | 0.991170 | 0.984255 |
| | **Thorax** | **Female** | 0.990890 | 0.994924 | 0.995122 | 0.883165 | 0.905518 | 0.906166 |
| | | **Male** | 0.996737 | 0.939085 | 0.925254 | 0.969958 | 0.960632 | 0.964784 |
| | **Viscera** | **Female** | 0.984629 | 0.980794 | 0.993663 | 0.919146 | 0.959881 | 0.971992 |
| | | **Male** | 0.989315 | 0.799956 | 0.751725 | 0.992267 | 0.992102 | 0.805981 |
| | **Abdomen** | **Female** | 0.993592 | 0.995429 | 0.996583 | 0.968139 | 0.980062 | 0.980141 |
| | | **Male** | 0.963463 | 0.936113 | 0.942848 | 0.979312 | 0.939376 | 0.927537 |
| | **Gonad** | **Female** | 0.994961 | 0.996948 | 0.993684 | 0.987874 | 0.975705 | 0.989799 |
| | | **Male** | 0.994361 | 0.994777 | 0.994190 | 0.973175 | 0.964849 | 0.982665 |
| **Dana** | **Head** | **Female** | 0.990367 | 0.989508 | 0.967599 | 0.994900 | 0.979750 | 0.997848 |
| | | **Male** | 0.990991 | 0.992224 | 0.993568 | 0.982729 | 0.987931 | 0.995473 |
| | **Thorax** | **Female** | 0.941168 | 0.994460 | 0.957395 | 0.987055 | 0.960592 | 0.997124 |
| | | **Male** | 0.944573 | 0.963127 | 0.994388 | 0.907228 | 0.987476 | 0.983303 |
| | **Viscera** | **Female** | 0.962710 | 0.932637 | 0.982137 | 0.986278 | 0.975557 | 0.963491 |
| | | **Male** | 0.993044 | 0.990088 | 0.991979 | 0.979297 | 0.988458 | 0.983946 |
| | **Abdomen** | **Female** | 0.991240 | 0.978971 | 0.959236 | 0.983975 | 0.967665 | 0.996980 |
| | | **Male** | 0.971883 | 0.964883 | 0.993465 | 0.965458 | 0.989691 | 0.990525 |
| | **Gonad** | **Female** | 0.983187 | 0.959404 | 0.988990 | 0.977506 | 0.995813 | 0.990288 |
| | | **Male** | 0.985149 | 0.978059 | 0.997103 | 0.991309 | 0.991224 | 0.990138 |
| **Dpse** | **Head** | **Female** | 0.993112 | 0.943349 | 0.946848 | 0.838438 | 0.840575 | 0.967611 |
| | | **Male** | 0.995843 | 0.989104 | 0.994996 | 0.985457 | 0.986487 | 0.987875 |
| | **Thorax** | **Female** | 0.954700 | 0.986829 | 0.948448 | 0.948178 | 0.898844 | 0.974722 |
| | | **Male** | 0.909400 | 0.909106 | 0.979517 | 0.931304 | 0.965471 | 0.969429 |
| | **Viscera** | **Female** | 0.991218 | 0.996787 | 0.984053 | 0.989554 | 0.979395 | 0.992060 |
| | | **Male** | 0.982862 | 0.980371 | 0.974773 | 0.918769 | 0.859093 | 0.907442 |
| | **Abdomen** | **Female** | 0.970571 | 0.979348 | 0.921108 | 0.952402 | 0.873621 | 0.992262 |
| | | **Male** | 0.771104 | 0.840642 | 0.802293 | 0.934902 | 0.758733 | 0.844536 |
| | **Gonad** | **Female** | 0.995431 | 0.982422 | 0.971455 | 0.928933 | 0.917062 | 0.962804 |
| | | **Male** | 0.996609 | 0.998662 | 0.996173 | 0.997104 | 0.993257 | 0.997435 |
| **Dper** | **Head** | **Female** | 0.984080 | 0.982618 | 0.960732 | 0.986538 | 0.988135 | 0.978817 |
| | | **Male** | 0.979499 | 0.988659 | 0.992045 | 0.980987 | 0.990617 | 0.988123 |
| | **Thorax** | **Female** | 0.975192 | 0.973009 | 0.960853 | 0.974363 | 0.974062 | 0.967320 |
| | | **Male** | 0.935970 | 0.864674 | 0.958101 | 0.908037 | 0.980721 | 0.950267 |
| | **Viscera** | **Female** | 0.995835 | 0.985633 | 0.986375 | 0.993985 | 0.993464 | 0.990415 |
| | | **Male** | 0.979966 | 0.956007 | 0.977611 | 0.988133 | 0.995732 | 0.974185 |
| | **Abdomen** | **Female** | 0.989332 | 0.958943 | 0.977370 | 0.959812 | 0.974976 | 0.960716 |
| | | **Male** | 0.965142 | 0.970291 | 0.970791 | 0.964362 | 0.995695 | 0.957293 |

Continued on next page

**Table S1 – continued from previous page**

| Species | Body part | Sex | R1vR2 | R1vR3 | R1vR4 | R2vR3 | R2vR4 | R3vR4 |
|---|---|---|---|---|---|---|---|---|
| | **Gonad** | **Female** | 0.961425 | 0.981003 | 0.990609 | 0.970611 | 0.995859 | 0.990006 |
| | | **Male** | 0.979448 | 0.935380 | 0.918542 | 0.985041 | 0.993212 | 0.934277 |
| **Dwil** | **Head** | **Female** | 0.997698 | 0.979226 | 0.978166 | 0.976718 | 0.976427 | 0.982758 |
| | | **Male** | 0.997537 | 0.997281 | 0.997059 | 0.997151 | 0.998717 | 0.998613 |
| | **Thorax** | **Female** | 0.982212 | 0.968952 | 0.968229 | 0.860930 | 0.878991 | 0.917237 |
| | | **Male** | 0.991469 | 0.993756 | 0.979984 | 0.990890 | 0.992521 | 0.981754 |
| | **Viscera** | **Female** | 0.945545 | 0.991329 | 0.937209 | 0.973417 | 0.928403 | 0.981598 |
| | | **Male** | 0.975367 | 0.989616 | 0.987774 | 0.969765 | 0.994070 | 0.990410 |
| | **Abdomen** | **Female** | 0.994095 | 0.978959 | 0.978260 | 0.963703 | 0.958889 | 0.949065 |
| | | **Male** | 0.985270 | 0.984225 | 0.978118 | 0.962820 | 0.956170 | 0.973740 |
| | **Gonad** | **Female** | 0.986676 | 0.969084 | 0.993086 | 0.975491 | 0.940041 | 0.907877 |
| | | **Male** | 0.997202 | 0.998340 | 0.996234 | 0.996643 | 0.991342 | 0.996868 |
| **Dvir** | **Head** | **Female** | 0.998306 | 0.954397 | 0.956706 | 0.988088 | 0.990246 | 0.921360 |
| | | **Male** | 0.995746 | 0.980533 | 0.979635 | 0.988538 | 0.986982 | 0.992969 |
| | **Thorax** | **Female** | 0.977790 | 0.993576 | 0.981624 | 0.993512 | 0.976819 | 0.990958 |
| | | **Male** | 0.992085 | 0.994168 | 0.987029 | 0.994137 | 0.995288 | 0.990043 |
| | **Viscera** | **Female** | 0.997040 | 0.990264 | 0.988539 | 0.977953 | 0.981485 | 0.981438 |
| | | **Male** | 0.985694 | 0.981124 | 0.986304 | 0.981461 | 0.991149 | 0.990728 |
| | **Abdomen** | **Female** | 0.991785 | 0.995693 | 0.998636 | 0.997905 | 0.993871 | 0.997094 |
| | | **Male** | 0.992116 | 0.994619 | 0.982653 | 0.991610 | 0.993747 | 0.983224 |
| | **Gonad** | **Female** | 0.991769 | 0.987540 | 0.988091 | 0.972785 | 0.978105 | 0.991981 |
| | | **Male** | 0.993468 | 0.995883 | 0.998775 | 0.998449 | 0.996464 | 0.997551 |
| **Dmoj** | **Head** | **Female** | 0.962843 | 0.960825 | 0.994306 | 0.943728 | 0.994027 | 0.992354 |
| | | **Male** | 0.989710 | 0.988592 | 0.993848 | 0.991283 | 0.994582 | 0.991813 |
| | **Thorax** | **Female** | 0.961575 | 0.963535 | 0.992263 | 0.959057 | 0.984834 | 0.988811 |
| | | **Male** | 0.958826 | 0.950014 | 0.874381 | 0.987367 | 0.924431 | 0.971886 |
| | **Viscera** | **Female** | 0.979529 | 0.982159 | 0.986241 | 0.957079 | 0.975849 | 0.974537 |
| | | **Male** | 0.987829 | 0.937080 | 0.941667 | 0.928200 | 0.933017 | 0.985201 |
| | **Abdomen** | **Female** | 0.931998 | 0.950575 | 0.981828 | 0.951998 | 0.995354 | 0.983527 |
| | | **Male** | 0.924865 | 0.966528 | 0.929062 | 0.980075 | 0.937620 | 0.977476 |
| | **Gonad** | **Female** | 0.972749 | 0.991417 | 0.986517 | 0.940641 | 0.988575 | 0.956016 |
| | | **Male** | 0.970933 | 0.982022 | 0.996073 | 0.987983 | 0.992238 | 0.996666 |
| **Dgri** | **Head** | **Female** | 0.974102 | 0.986001 | 0.942319 | 0.993673 | 0.975433 | 0.984177 |
| | | **Male** | 0.996217 | 0.979935 | 0.980031 | 0.985497 | 0.992074 | 0.966464 |
| | **Thorax** | **Female** | 0.979418 | 0.982102 | 0.951328 | 0.984850 | 0.971397 | 0.981054 |
| | | **Male** | 0.996772 | 0.991701 | 0.993935 | 0.992323 | 0.993478 | 0.993230 |
| | **Viscera** | **Female** | 0.950148 | 0.976583 | 0.966260 | 0.983401 | 0.970601 | 0.984913 |
| | | **Male** | 0.972196 | 0.963684 | 0.987954 | 0.941835 | 0.978413 | 0.984942 |
| | **Abdomen** | **Female** | 0.960385 | 0.929732 | 0.914441 | 0.854547 | 0.749440 | 0.759291 |
| | | **Male** | 0.931437 | 0.940168 | 0.927029 | 0.925515 | 0.900744 | 0.896803 |
| | **Gonad** | **Female** | 0.846158 | 0.879609 | 0.682405 | 0.978817 | 0.742362 | 0.880504 |
| | | **Male** | 0.991673 | 0.915842 | 0.931773 | 0.990275 | 0.988463 | 0.906694 |

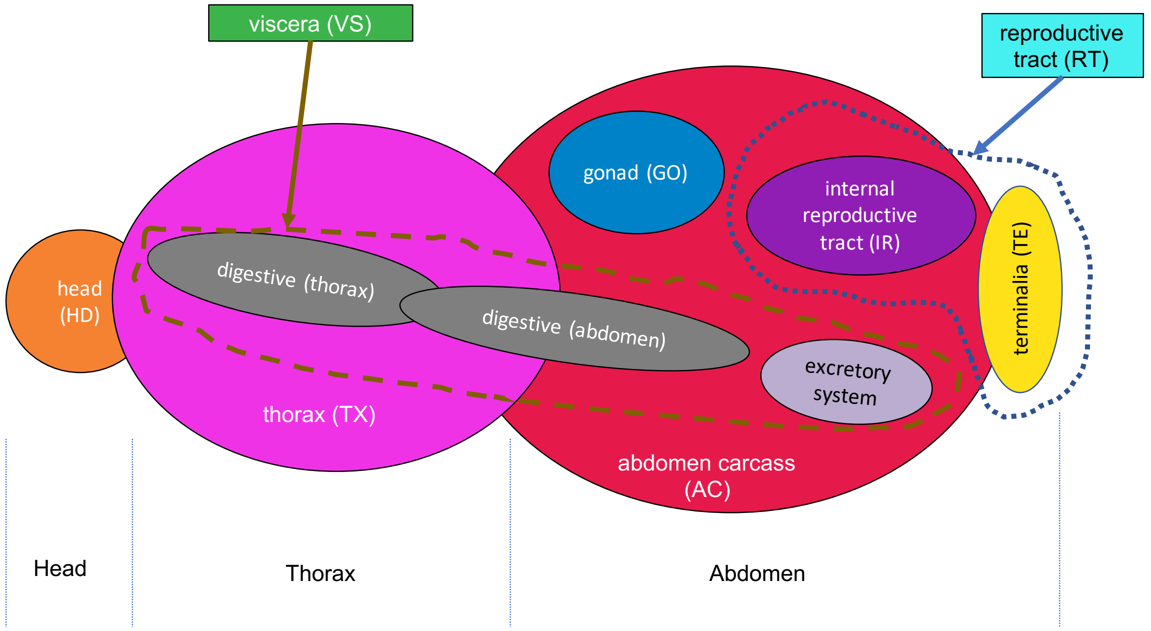


Figure S2: **An illustrative cartoon for the body parts of a fruit-fly from where the samples were taken and bulk RNA-seq data was generated.** Only the following five body parts were used for our analysis as only they all had all 4 replicates: head (HD), thorax (TX), viscera (VS), abdomen carcass (AC) and gonad (GO).

##### S2.2 Principal component analysis

Figure S5 shows the PCA clustering obtained from the average expression of all replicates. The PCA was done using scikit-learn library in Python. The similarity measure was 1-Spearman's correlation coefficient. The analysis confirms clustering by body parts.

##### S2.3 Hierarchical clustering

Hierarchical clustering was obtained by UPGMA method (R function `agnes` method `average`). Figure S6a (repeated from Fig. 3a in main text) shows the clustering obtained when the distance given by 1-Spearman's correlation coefficient on the normalized gene expression values. We observe that the expression data predominantly clusters by tissue. However, when we used log gene expression values and Euclidean distances then related tissues of the same species often clustered first (Figure S6b). Specifically, with this similarity measure, head and thorax typically cluster together. This reflects the fact that in fly these two body parts are related. Thus both tissue-specific and species-specific trends can be observed, depending on the weight given to genes in the tails of the expression distributions.

#### S3 EvoGeneX

EvoGeneX takes as its input i) a rooted evolutionary tree, ii) the values of quantitative characters (with biological replicates) for all terminal taxa, and iii) the labelling of all nodes for which regime

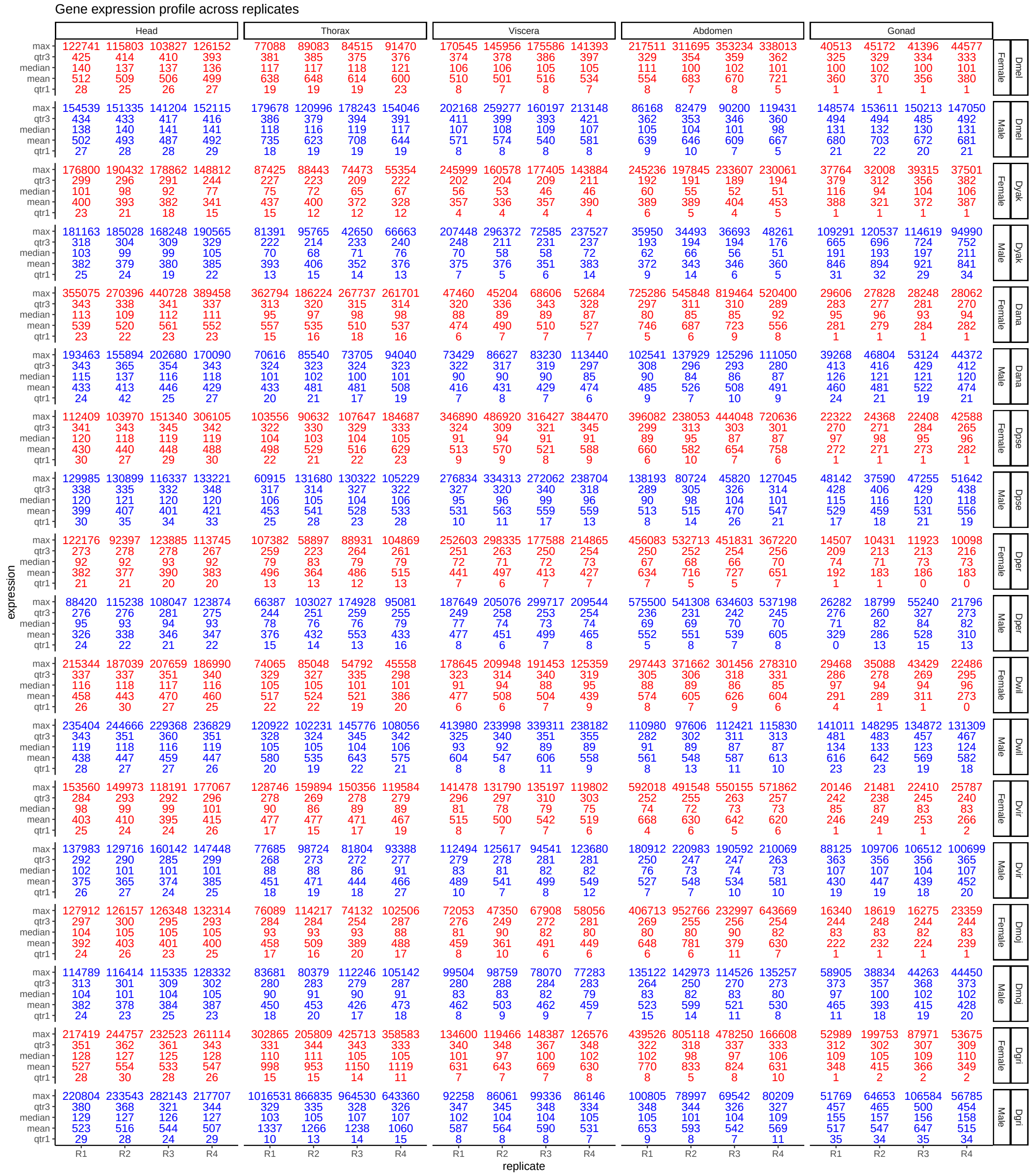


Figure S3: **Summary statistics of the replicate expression across species, body parts and sex.**

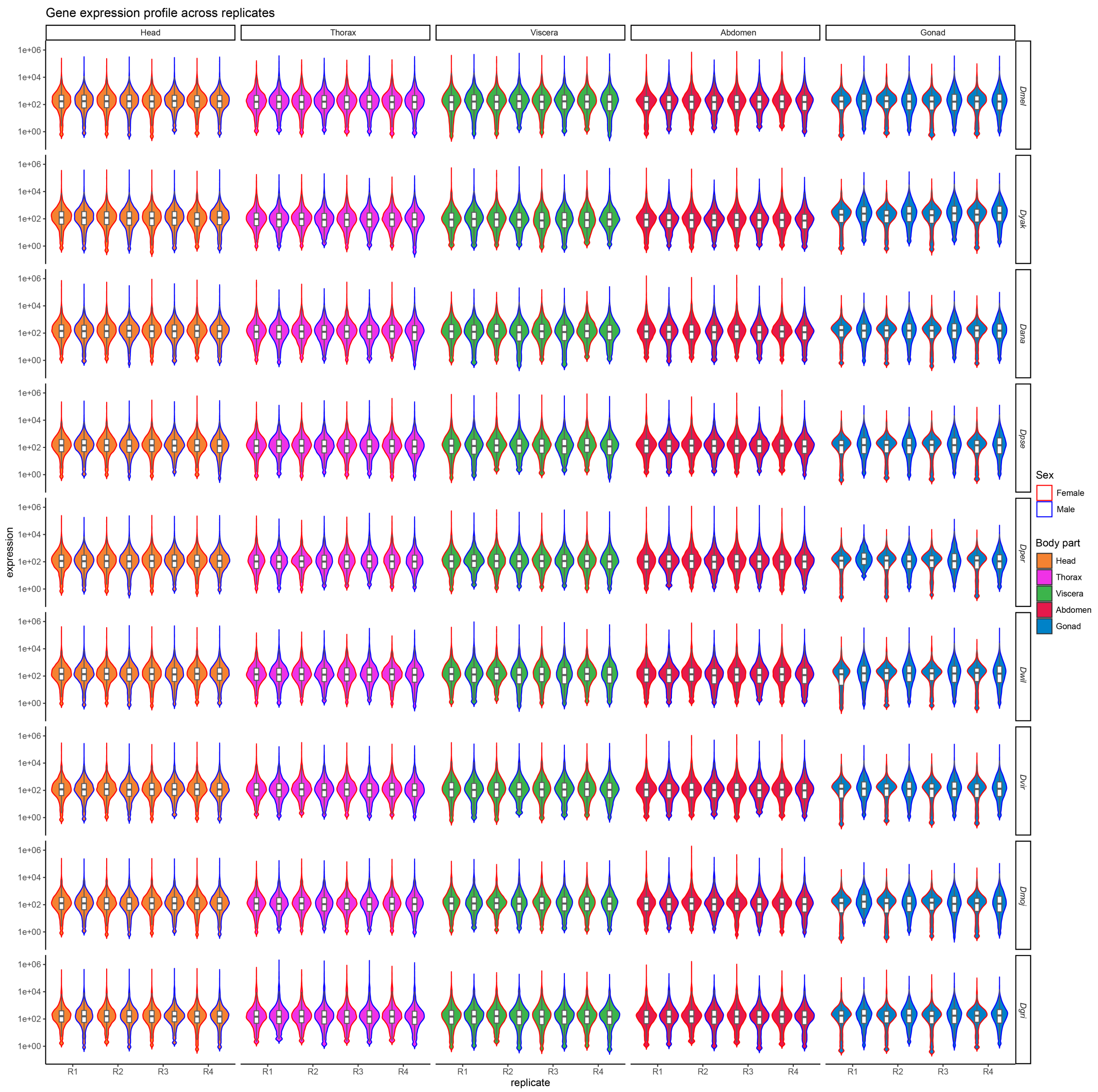


Figure S4: **Violin-plots of the replicate expression across species, body parts and sex.**

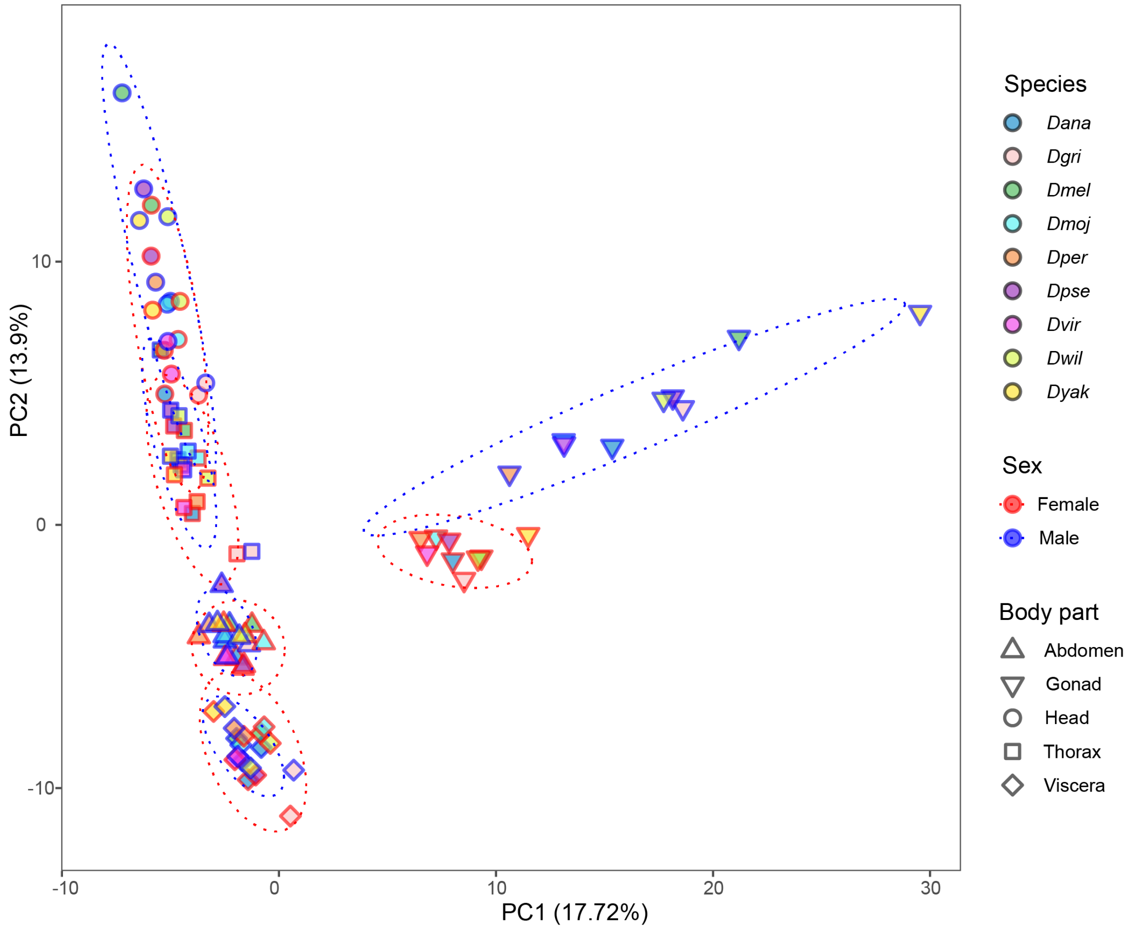


Figure S5: **Principal Component Analysis of gene expression data (replicate mean).** The gonad expressions cluster by sex first and then by species whereas non-gonad expressions cluster by body parts first and then by species and sex, in that order.

it belongs. EvoGeneX estimates i) the parameters associated with three models, and reports ii) optimum Loglikehood value.

##### S3.1 Inference of the expected value and covariance in the EvoGeneX model – complete derivations

Specifically, let $T$ be a rooted evolutionary tree with $N$ terminal taxa (here species) where the branch lengths represent evolutionary time, and the root represents the last common ancestor (LCA) of all $N$ taxa. For each specie $i$, the path from the root to the corresponding terminal node represents its lineage. Let $T_i$ be the total length of lineage $i$ and let $X_i(t)$ denote the value of the quantitative character of interest (here gene expression) for the $i$th lineage at time $t$. We assume $t = 0$ at the root of the tree. For two species $i, j$ let $s_{i,j}$ denote the time of the speciation event when lineages $i$ and $j$ diverged, and consequently, $X_i(t) = X_j(t)$ for all $t < s_{i,j}$.

We assume that for each lineage $i$, $X_i(t)$ evolves according to a mean-reverting Ornstein-

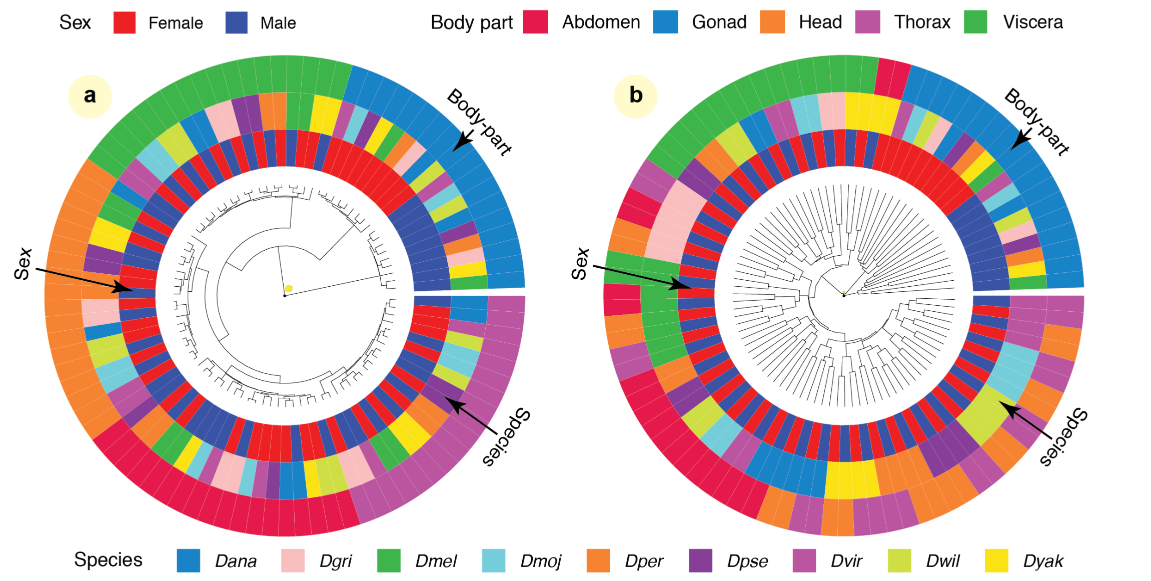


Figure S6: **Hierarchical clustering of gene expression data using two different distance matrices.** The outside circles correspond to body part, the middle circles to species and the inside circles to sex. (**a**) Using Spearman's correlation: The main organizing factor is cluster first (except for gonads which cluster first by sex). (**b**) Using Euclidean distance: Here, related tissues, e.g., head, thorax, and abdomen carcass, for the same species often cluster together. This emphasizes the importance of choosing distance measures for inferences based on clustering of expressions.

Uhlenbeck (OU) process (Hansen 1997)

$$dX_i(t) = \alpha\left[\beta_i(t) - X_i(t)\right]dt + \sigma dB_i(t) \tag{S1}$$

where $0 \leq t \leq T$, $\alpha$ is the rate of mean reversion (the strength of the pull to the optimum), $\sigma$ is the volatility of the process, and the function $\beta_i(t)$ represents the dynamic optimum trait value of the OU process and as such identifies the selection regime acting on lineage $i$ over the course of its history. $dB_i(t)$ denotes increments of a standard Brownian motion (BM); heuristically, we can think of them as normal random variables with mean 0 and variance $dt$.

Multiplying both sides of (S1) by $e^{\alpha t}$ we get

$$dX_i(t)e^{\alpha t} = \alpha e^{\alpha t}\beta_i(t)dt - \alpha e^{\alpha t}X_i(t)dt + \sigma e^{\alpha t}dB_i(t)$$

or

$$dX_i(t)e^{\alpha t} + \alpha e^{\alpha t}X_i(t)dt = \alpha e^{\alpha t}\beta_i(t)dt + \sigma e^{\alpha t}dB_i(t)$$

implying

$$d(X_i(t)e^{\alpha t}) = \alpha e^{\alpha t}\beta_i(t)dt + \sigma e^{\alpha t}dB_i(t)$$

Integrating both sides from $t = 0$ to $t = t$ we get

$$X_i(t)e^{\alpha t} - X_i(0) = \int_0^t \alpha e^{\alpha s}\beta_i(s)ds + \int_0^t \sigma e^{\alpha s} dB_i(s)$$

implying

$$X_i(t) = e^{-\alpha t}X_i(0) + e^{-\alpha t}\int_0^t \alpha e^{\alpha s}\beta_i(s)ds + e^{-\alpha t}\int_0^t \sigma e^{\alpha s} dB_i(s)$$

Because this defines a Gaussian process, the first moment of $X_i(t)$ is completely specified by the deterministic (or non-stochastic) components as the stochastic component has mean 0. In particular, we have the expected trait value for species $i$ as

$$\mathrm{E}[X_i(t) \mid X_i(0) = \theta_0] = \theta_0 e^{-\alpha t} + e^{-\alpha t}\int_0^t \alpha e^{\alpha s}\beta_i(s)ds$$

and the covariance between species $i$ and species $j$ is

$$\begin{aligned}
&\mathrm{Cov}[X_i(t_i), X_j(t_j) \mid X_i(0) = X_j(0) = \theta_0] \\
&= \mathrm{E}[(X_i(t_i) - E[X_i(t_i)] \mid X_i(0) = \theta_0)(X_j(t_j) - E[X_j(t_j)] \mid X_j(0) = \theta_0)] \\
&= \mathrm{E}\left[\left(e^{-\alpha t_i}\int_0^{t_i} \sigma e^{\alpha u} dB_i(u)\right)\left(e^{-\alpha t_j}\int_0^{t_j} \sigma e^{\alpha v} dB_j(v)\right)\right] \\
&= \mathrm{E}\left[\sigma^2 e^{-\alpha(t_i+t_j)}\left(\int_0^{t_i} e^{\alpha u} dB_i(u)\right)\left(\int_0^{t_j} e^{\alpha v} dB_j(v)\right)\right] \\
&= \sigma^2 e^{-\alpha(t_i+t_j)}\mathrm{E}\left[\left(\int_0^{s_{i,j}} e^{\alpha u} dB_i(u) + \int_{s_{i,j}}^{t_i} e^{\alpha u} dB_i(u)\right)\left(\int_0^{s_{i,j}} e^{\alpha v} dB_j(v) + \int_{s_{i,j}}^{t_j} e^{\alpha v} dB_j(v)\right)\right] \\
&= \sigma^2 e^{-\alpha(t_i+t_j)}\mathrm{E}\left[\begin{array}{l}
\underbrace{\left(\int_0^{s_{i,j}} e^{\alpha u} dB_i(u)\right)\left(\int_0^{s_{i,j}} e^{\alpha v} dB_j(v)\right)}_{\boldsymbol{A}} \\
+\underbrace{\left(\int_0^{s_{i,j}} e^{\alpha u} dB_i(u)\right)\left(\int_{s_{i,j}}^{t_j} e^{\alpha v} dB_j(v)\right)}_{\boldsymbol{B}} \\
+\underbrace{\left(\int_{s_{i,j}}^{t_i} e^{\alpha u} dB_i(u)\right)\left(\int_0^{s_{i,j}} e^{\alpha v} dB_j(v)\right)}_{\boldsymbol{C}} \\
+\underbrace{\left(\int_{s_{i,j}}^{t_i} e^{\alpha u} dB_i(u)\right)\left(\int_{s_{i,j}}^{t_j} e^{\alpha v} dB_j(v)\right)}_{\boldsymbol{D}}
\end{array}\right] \\
&= \sigma^2 e^{-\alpha(t_i+t_j)}\mathrm{E}\left[\underbrace{\left(\int_0^{s_{i,j}} e^{\alpha u} dB_i(u)\right)\left(\int_0^{s_{i,j}} e^{\alpha v} dB_i(v)\right)}_{A}\right] \quad \text{(by independent increment property of Brownian motion, each of } \boldsymbol{B, C, D} \text{ is 0 and } B_i(t) = B_j(t) \text{ for } t < s_{i,j}) \\
&= \sigma^2 e^{-\alpha(t_i+t_j)}\mathrm{E}\left[\int_0^{s_{i,j}} e^{2\alpha u} du\right] \quad \text{(by Itô Isometry on } \boldsymbol{A})
\end{aligned}$$

$$= \sigma^2 e^{-\alpha(t_i+t_j)} \mathrm{E}\left[\frac{1}{2\alpha}\left(e^{2\alpha s_{i,j}} - 1\right)\right] = \frac{\sigma^2}{2\alpha} e^{-\alpha(t_i+t_j)}(e^{2\alpha s_{i,j}} - 1) = \frac{\sigma^2}{2\alpha} e^{-\alpha(t_i+t_j-2s_{i,j})}(1 - e^{-2\alpha s_{i,j}})$$

Given multiple observations $Y_{i,k}$ for the trait value $X_i(T_i)$ for the terminal taxa $i$, the observed variance (including technical, environmental) could be explained by within-species variance ($\gamma^2$). Thus the trait value of $k$th replicate of species $i$ is

$$Y_{i,k}(t) = X_i(t) + \varepsilon_{i,k} \tag{S2}$$

where each $\varepsilon_{i,k} \sim N(0, \gamma\sigma^2)$ is independent and identically distributed. Thus, we have the expected trait value for replicate $k$ of species $i$ as

$$\mathrm{E}[Y_{i,k}(t_i) \mid X_i(0) = \theta_0] = \mathrm{E}[X_i(t_i) \mid X_i(0) = \theta_0] = \theta_0 e^{-\alpha t_i} + e^{-\alpha t_i} \int_0^{t_i} \alpha e^{\alpha s} \beta_i(s) ds \tag{S3}$$

and the covariance between species $i$, replicate $k$ and species $j$, replicate $l$ is

$$\begin{aligned}
&\mathrm{Cov}[Y_{i,k}(t_i), Y_{j,l}(t_j) \mid X_i(0) = X_j(0) = \theta_0] \\
&= \mathrm{Cov}[X_i(t_i), X_j(t_j) \mid X_i(0) = X_j(0) = \theta_0] + \begin{cases} \gamma\sigma^2 & \text{if } (i = j) \text{ and } (k = l) \\ 0 & \text{otherwise} \end{cases} \\
&= \frac{\sigma^2}{2\alpha} e^{-\alpha(t_i+t_j-2s_{i,j})}(1 - e^{-2\alpha s_{i,j}}) + \begin{cases} \gamma\sigma^2 & \text{if } (i = j) \text{ and } (k = l) \\ 0 & \text{otherwise} \end{cases}
\end{aligned} \tag{S4}$$

##### S3.2 Different choices for the optimum-defining function $\beta_i(t)$, describe different modes of evolution – complete derivations

Equations (S3) and (S4) providing the expectation and covariance of different observations at the taxa are very general. Using different definitions of $\beta_i(t)$ leads to various biologically relevant models. The simplest model correspond to the situation where, for all $i$, $\beta_i(t)$ is a constant. This describes the *constrained evolution* model corresponding to one common optimum value.

Under the assumption of multiple optima, it is reasonable to assume that $\beta_i(t)$ changes at speciation events and remains constant along individual edges of the phylogenic tree. We call the times of speciation events "epochs." The history of the $i$th lineage consists of a number, $\kappa(i)$, of sequential branch segments demarcated by epochs $0 = t_i^0 < t_i^1 < t_i^2 < \ldots < t_i^{\kappa(i)} = T_i$. Thus, for $t_i^{\tau-1} \le t \le t_i^\tau$ equation (S3) takes the following form

$$\begin{aligned}
\mathrm{E}[Y_{i,k}(T_i) \mid X_i(0) = \theta_0] &= \theta_0 e^{-\alpha T_i} + e^{-\alpha T_i} \int_0^{T_i} \alpha e^{\alpha s} \beta_i(s) ds \\
&= \theta_0 e^{-\alpha T_i} + e^{-\alpha T_i} \sum_{\tau=1}^{\kappa(i)} \int_{t_i^{\tau-1}}^{t_i^\tau} \alpha e^{\alpha s} \beta_i^\tau ds \\
&= \theta_0 e^{-\alpha T_i} + e^{-\alpha T_i} \sum_{\tau=1}^{\kappa(i)} \left(e^{\alpha t_i^\tau} - e^{\alpha t_i^{\tau-1}}\right) \beta_i^\tau \\
&= \theta_0 e^{-\alpha T_i} + \sum_{\tau=1}^{\kappa(i)} \left(e^{-\alpha(T_i - t_i^\tau)} - e^{-\alpha(T_i - t_i^{\tau-1})}\right) \beta_i^\tau .
\end{aligned} \tag{S5}$$

The number of distinct $\beta_i^\tau$ might be as big as the number of tree edges $(2N - 1)$. However, it is not practical to estimate so many parameters. Instead, following Hansen (1997), we assume a small number, $R$, of distinct optimum values $\theta_r$, $r = 1, \ldots, R$, each corresponding to one *selective regime*. In fact one of the most interesting cases corresponds to the model with two optima where one branch of the tree follows a regime of optimum values $\theta_1$ and the rest of the tree $\theta_0$ (J. Chen et al. 2019; Brawand et al. 2011).

Let the binary variable $\beta_{i,r}^{\tau}$ represent if the $\tau$th branch on lineage $i$ has operated in $r$th regime. Then we have $\beta_i^{\tau} = \sum_{r=1}^{R} \beta_{i,r}^{\tau}\theta_r$ and rearranging similar terms in equation (S5) gives

$$\begin{aligned} \mathrm{E}[Y_{i,k}(T_i) \mid X_i(0) = \theta_0] &= \theta_0 e^{-\alpha T_i} + \sum_{\tau=1}^{\kappa(i)} \left(e^{-\alpha(T_i - t_i^{\tau})} - e^{-\alpha(T_i - t_i^{\tau-1})}\right) \sum_{r=1}^{R} \beta_{i,r}^{\tau}\theta_k \\ &= \theta_0 e^{-\alpha T_i} + \sum_{r=1}^{R} \left( \sum_{\tau=1}^{\kappa(i)} \left(e^{-\alpha(T_i - t_i^{\tau})} - e^{-\alpha(T_i - t_i^{\tau-1})}\right) \beta_{i,r}^{\tau} \right) \theta_r \end{aligned} \tag{S6}$$

Note that since each branch is associated with exactly one optimum, for each $i, \tau$ there is exactly one index $r$ such that $\beta_{i,r}^{\tau} = 1$ and $\beta_{i,r'}^{\tau} = 0$ for all $r \neq r'$. Further, self-consistency requires that $\beta_{i,r}^{\tau} = \beta_{j,r}^{\eta}$ whenever lineage $i$ and $j$ share the branch ending in epoch $t_i^{\tau} = t_j^{\eta}$.

The parameters $\theta_0, \theta_1, \ldots, \theta_R$, together with $\alpha, \sigma, \gamma$ must be estimated which we do using commonly used Maximum Likelihood (ML) estimation method.

#### S3.3 Maximum likelihood estimates – complete derivations

In the following, it will be convenient to make use of matrix notation. Accordingly, we collect our random variables, $X_i(t)$ for the trait values at the taxa, and $Y_{i,k}(t)$ for the replicated trait values, in the vectors $\mathbf{x}(t)$ and $\mathbf{y}(t)$, respectively, and our observed quantitative data in the vector $\mathbf{y}$ with components $y_{i+(k-1)N} = Y_{i,k}(T_i)$, the observed value of the quantitative character in each species $i$, $i = 1, \ldots, N$, at the end of an evolutionary process of length time $= T_i$. Thus, equation (S6) can be rewritten in matrix notation as

$$\mathrm{E}[\mathbf{y}(T) \mid \mathbf{x}(0) = \theta_0 \mathbb{1}] = \mathbf{W}\boldsymbol{\theta} \tag{S7}$$

where $\mathbb{1}$ is a vector of all 1s, column vector $\boldsymbol{\theta} = (\theta_0, \theta_1, \ldots, \theta_R)^T$ and the weight matrix $W$ is dependent only on $\alpha$ among the parameters and has entries

$$\begin{aligned} W_{i+(k-1)N,0} &= e^{-\alpha T_i} \\ W_{i+(k-1)N,r} &= \sum_{\tau=1}^{\kappa(i)} \left(e^{-\alpha(T_i - t_i^{\tau})} - e^{-\alpha(T_i - t_i^{\tau-1})}\right) \beta_{i,r}^{\tau} \end{aligned} \tag{S8}$$

for $i = 1, \ldots, N$, $k = 1, \ldots, M$ and $r = 1, \ldots, R$. Similarly, let an $MN \times MN$ matrix $\mathbf{V}$ denote the covariance matrix where the covariance between species $i$, replicate $k$ and species $j$, replicate $l$ is given by the entry

$$\begin{aligned} v_{(i,k),(j,l)} = v_{i+(k-1)N, j+(l-1)N} &= \mathrm{Cov}[Y_{i,k}(t_i), Y_{j,l}(t_j) \mid X_i(0) = X_j(0) = \theta_0] \\ &= \frac{\sigma^2}{2\alpha} e^{-\alpha(t_i + t_j - 2s_{i,j})}(1 - e^{-2\alpha s_{i,j}}) + \begin{cases} \gamma\sigma^2 & \text{if } (i = j) \text{ and } (k = l) \\ 0 & \text{otherwise.} \end{cases} \end{aligned} \tag{S9}$$

It is known that $\mathbf{y}$ follows a multi-variate Gaussian distribution $\mathcal{N}(\mathbf{W}\boldsymbol{\theta}, \mathbf{V})$ with mean and co-variance given by equations (S7) and (S9) (Hansen and Martins 1996). Thus, the likelihood of the parameters $\alpha, \sigma, \gamma$, and $\boldsymbol{\theta}$, given the data $\mathbf{y}(T) = \mathbf{y}$, is

$$\mathcal{L}(\alpha, \sigma, \gamma, \boldsymbol{\theta} \mid \mathbf{y}) = \frac{1}{\sqrt{(2\pi)^{NM} \det \mathbf{V}}} \exp\left[-\frac{(\mathbf{y} - \mathbf{W}\boldsymbol{\theta})^T \mathbf{V}^{-1} (\mathbf{y} - \mathbf{W}\boldsymbol{\theta})}{2}\right] \tag{S10}$$

As maximizing $\mathcal{L}$ is equivalent to minimizing $U = -2\log\mathcal{L}$, we seek to minimize

$$U(\alpha, \sigma, \gamma, \boldsymbol{\theta} \mid \mathbf{y}) = NM\log(2\pi) + \log\det\mathbf{V} + (\mathbf{y} - \mathbf{W}\boldsymbol{\theta})^T\mathbf{V}^{-1}(\mathbf{y} - \mathbf{W}\boldsymbol{\theta}) \tag{S11}$$

However, it can be noted that $\mathbf{V}$ has a nice structure and can be expressed as $\sigma^2(\tilde{\mathbf{V}} + \gamma\mathbb{I})$ where $\tilde{\mathbf{V}}$ is dependent on $\alpha$ only among all the parameters and $\mathbb{I}$ is an identity matrix of size $MN$. The elements of $\tilde{\mathbf{V}}$ are given by $\tilde{V}_{(i,k),(j,l)} = \frac{1}{2\alpha}e^{-\alpha(t_i+t_j-2s_{i,j})}(1 - e^{-2\alpha s_{i,j}})$.

Thus, $U$ can be expressed as

$$U(\alpha, \sigma, \gamma, \boldsymbol{\theta} \mid \mathbf{y}) = NM\log(2\pi\sigma^2) + \log\det(\tilde{\mathbf{V}} + \gamma\mathbb{I}) + \frac{1}{\sigma^2}(\mathbf{y} - \mathbf{W}\boldsymbol{\theta})^T(\tilde{\mathbf{V}} + \gamma\mathbb{I})^{-1}(\mathbf{y} - \mathbf{W}\boldsymbol{\theta})$$

whose minimum can be estimated using any off-the-self nonlinear optimization solver.

However, we improve the efficiency by utilizing Karush-Kuhn-Tucker conditions at the minimum solutions. Taking partial derivatives of $U$ with respect to $\sigma$ and $\boldsymbol{\theta}$ and equating them to 0 at an optimal solution $(\hat{\alpha}, \hat{\sigma}, \hat{\gamma}, \hat{\boldsymbol{\theta}})$, we get

$$\begin{aligned} \hat{\sigma}^2 &= \frac{1}{NM}(\mathbf{y} - \mathbf{W}\hat{\boldsymbol{\theta}})^T(\tilde{\mathbf{V}} + \hat{\gamma}\mathbb{I})^{-1}(\mathbf{y} - \mathbf{W}\hat{\boldsymbol{\theta}}), \text{ and} \\ \hat{\boldsymbol{\theta}} &= \left(\mathbf{W}^T(\tilde{\mathbf{V}} + \hat{\gamma}\mathbb{I})^{-1}\mathbf{W}\right)^{-1}\mathbf{W}^T(\tilde{\mathbf{V}} + \hat{\gamma}\mathbb{I})^{-1}\mathbf{y} \end{aligned} \tag{S12}$$

Thus, instead of minimizing function $U$ of four parameters $\alpha, \sigma, \gamma$ and $\boldsymbol{\theta}$, it is enough to minimize a new function $\tilde{U}$ of two parameters, $\alpha$ and $\gamma$,

$$\tilde{U}(\alpha, \gamma) = NM\left[1 + \log 2\pi\hat{\sigma}^2(\alpha, \gamma)\right] + \log\det(\tilde{\mathbf{V}} + \gamma\mathbb{I}) \tag{S13}$$

where the following two intermediate functions

$$\begin{aligned} \hat{\boldsymbol{\theta}}(\alpha, \gamma) &= \left(\mathbf{W}^T(\tilde{\mathbf{V}} + \gamma\mathbb{I})^{-1}\mathbf{W}\right)^{-1}\mathbf{W}^T(\tilde{\mathbf{V}} + \gamma\mathbb{I})^{-1}\mathbf{y} \\ \hat{\sigma}^2(\alpha, \gamma) &= \frac{1}{NM}(\mathbf{y} - \mathbf{W}\hat{\boldsymbol{\theta}}(\alpha, \gamma))^T(\tilde{\mathbf{V}} + \gamma\mathbb{I})^{-1}(\mathbf{y} - \mathbf{W}\hat{\boldsymbol{\theta}}(\alpha, \gamma)) \end{aligned} \tag{S14}$$

give the values of the remaining two parameters $\sigma$ and $\boldsymbol{\theta}$ at the optimal solution.

#### S3.4 ML estimates for Brownian model – complete derivations

We need to compute maximum likelihood estimate for the Brownian model as well for comparing with the EvoGeneX model of evolution. BM is a simplified model in comparison to OU: there is no “attracting” optimal values and hence there is no $\alpha$ parameter and $\theta$ has only one value to be estimated corresponding to $\theta_0$.

Starting with the stochastic equation $Y_{i,k}(t) = X_i(t) + \varepsilon_{i,k} \sim N(0, \gamma\sigma^2)$ and the stochastic differential equation $X_i(t) = \sigma dB_i(t)$ we can follow the steps in the previous sections to derive the following log likelihood function $U = -2\log\mathcal{L}$

$$U(\sigma, \gamma, \theta_0 \mid \mathbf{y}) = NM\log(2\pi\sigma^2) + \log\det(\tilde{\mathbf{V}} + \gamma\mathbb{I}) + \frac{1}{\sigma^2}(\mathbf{y} - \theta_0\mathbb{1})^T(\tilde{\mathbf{V}} + \gamma\mathbb{I})^{-1}(\mathbf{y} - \theta_0\mathbb{1}) \tag{S15}$$

where the elements of $\tilde{\mathbf{V}}$ are given by $\tilde{V}_{(i,k),(j,l)} = s_{i,j}$, the time from the root of the phylogenetic tree to the least common ancestor of the taxa $i, j$. Intuitively, under Brownian motion, changes in trait values over any interval of time $t$ are always drawn from a normal distribution with mean 0 and variance proportional to the product of the rate of evolution and the length of time ($\sigma^2 t$).

Using Karush-Kuhn-Tucker conditions at the minimum solutions, we can show that it is enough to solve a function $\tilde{U}(\gamma)$ of one parameter $\gamma$

$$\tilde{U}(\gamma) = NM\left[1 + \log 2\pi\hat{\sigma}^2(\gamma)\right] + \log\det(\tilde{\mathbf{V}} + \gamma\mathbb{I}) \tag{S16}$$

where the following two intermediate functions

$$\begin{aligned} \hat{\theta}_0(\gamma) &= \left(\mathbb{1}^T(\tilde{\mathbf{V}} + \gamma\mathbb{I})^{-1}\mathbb{1}\right)^{-1}\mathbb{1}^T(\tilde{\mathbf{V}} + \gamma\mathbb{I})^{-1}\mathbf{y} \\ \hat{\sigma}^2(\alpha, \gamma) &= \frac{1}{NM}(\mathbf{y} - \hat{\theta}_0(\gamma)\mathbb{1})^T(\tilde{\mathbf{V}} + \gamma\mathbb{I})^{-1}(\mathbf{y} - \hat{\theta}_0(\gamma)\mathbb{1}) \end{aligned} \tag{S17}$$

give the values of the remaining two parameters $\sigma$ and $\theta_0$ at the optimal solution.

##### S3.5 Computing statistical significance

We use statistical hypothesis testing to decide which of the three different modes of evolution the trait has undergone: i) neutral, ii) constrained and ii) adaptive (see Section 2.1 in main text). For this purpose we use likelihood ratio test. Specifically, for two models $H_0$, $H_1$ with parameters $\mathbf{\Theta}_0, \mathbf{\Theta}_1$ and likelihoods $L_0(\mathbf{\Theta}_0), L_1(\mathbf{\Theta}_1)$, to test if $H_1$ models the data better than $H_0$, we check if the ratio of likelihoods, $\lambda = L_1/L_0$ is significantly greater than 1. By a result of Wilks (1938), the test statistic $-2\log(\lambda)$ is asymptotically $\chi^2$ distributed degrees of freedom equal to the difference in dimensionality of $\mathbf{\Theta}_1$ and $\mathbf{\Theta}_0$. Thus the statistical significance is provided by the upper tail probability $\mathrm{P}[x > -2\log(\lambda)]$ of $\chi^2$ distribution.

#### S4 Simulations

Figure S7 extends Fig. 2a by incorporating the performance metrics of a version of OUCH (King and Butler 2009) that treats replicated expression values as multivariate trait whose components co-evolve in time. This version which we call OUCH.MV, needs optimization over a large number of parameters (in the order of squared number of replicates) and seems to work poorly in datasets of smaller number of species like ours. We also incorporate the $auPRC$ metrics for all three method if we relax the constraint on the parameter values that the within-species variance should not exceed the asymptotic between species variance (corresponding to the case $\gamma$ all instead of $\gamma$: valid).

Figure S8 extends Fig. 2 by also incorporating the performance metric for fold values less than 1. It shows that the performance of EvoGeneX is not affected if the optimum expression for a branch goes down instead of going up by the same amount of fold change.

#### S5 Genes with neutral expression reconstruct expression based phylogenetic tree close to the sequence based tree

We reconstructed the phylogenetic tree from the expression values of all genes across species for each body part and sex separately using the neighbour joining (NJ) method and the Spearman's correlation based distances. We repeated the same considering only the genes whose expression evolution could not be rejected as neutral by EvoGeneX. The Robinson-Foulds distances of the reconstructed trees from the known sequence based tree (Section S1.1) are shown in Table S2. It can be noted that the trees constructed using only the genes undergoing neutral (or nearly neutral) expression evolution are often closer to the sequence based evolutionary tree than the tree constructed using all genes.

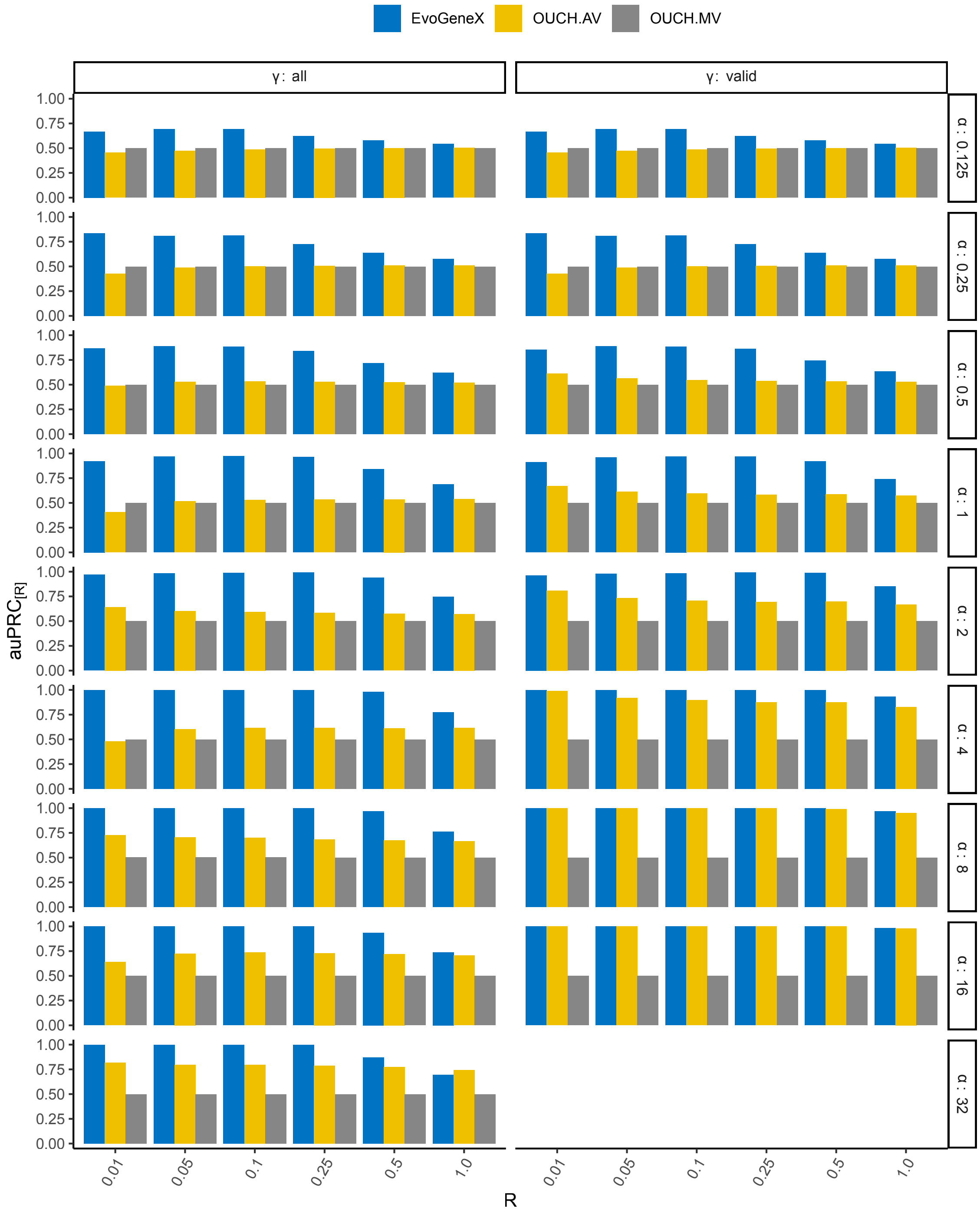


Figure S7: **Detailed results on single-regime simulations**

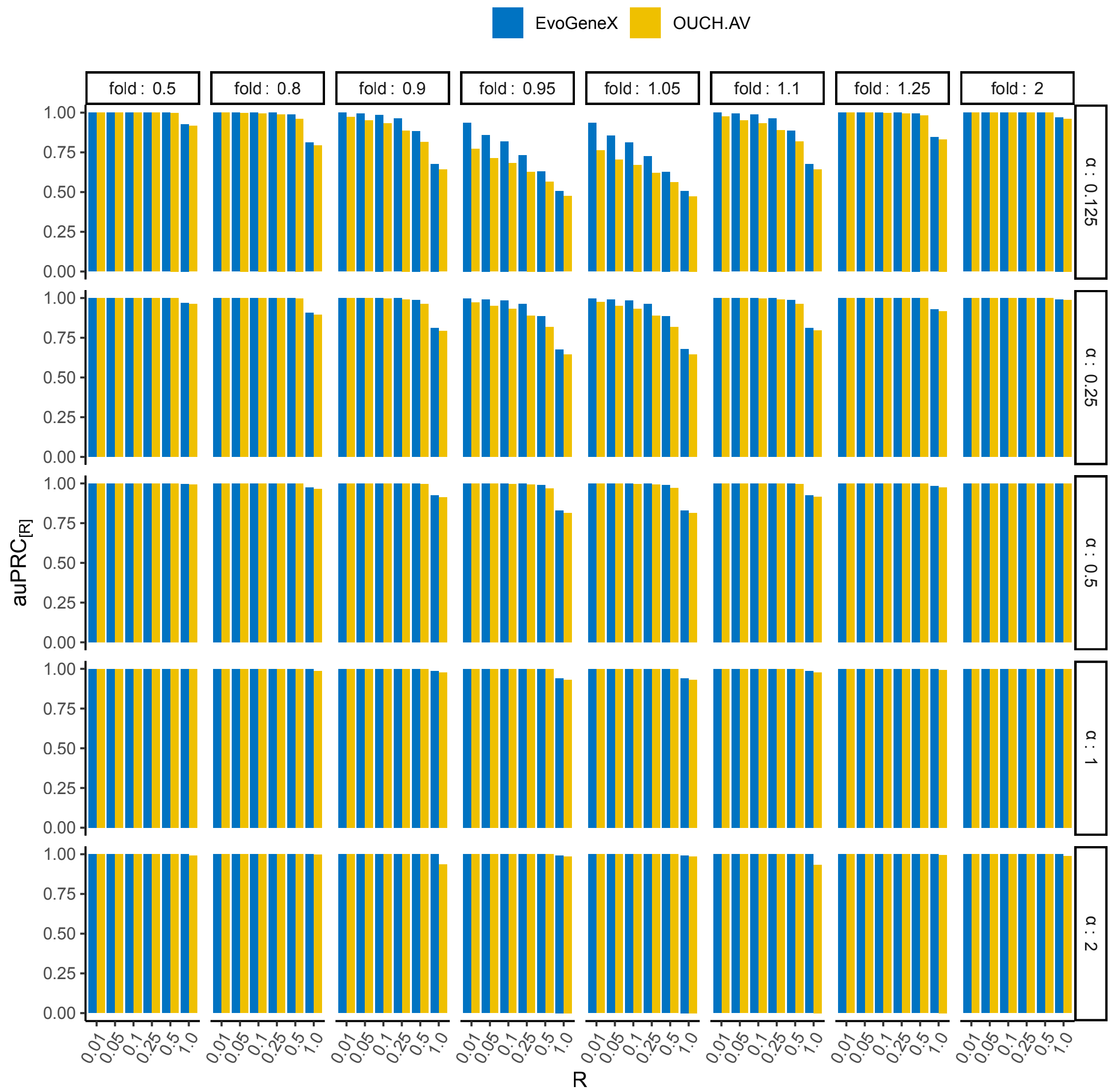


Figure S8: **Detailed results on two-regime simulations**

Table S2: **Robinson-Foulds distance of the known sequence based tree (Section S1.1) from the Neighbour Joining tree obtained by using Spearman's correlation based distance of the expression values considering three gene groups.** (i) all genes, (ii) neutrally evolved of genes as detected by EvoGeneX, and (iii) constrained genes.

| **Body part** | **Genes** **Sex** | All | Neutral | Constrained |
|---|---|---|---|---|
| **Head** | **Female** | 0 | 0 | 6 |
| | **Male** | 0 | 2 | 8 |
| **Thorax** | **Female** | 2 | 2 | 10 |
| | **Male** | 2 | 2 | 8 |
| **Viscera** | **Female** | 2 | 2 | 4 |
| | **Male** | 2 | 0 | 4 |
| **Abdomen** | **Female** | 2 | 0 | 6 |
| | **Male** | 0 | 0 | 4 |
| **Gonad** | **Female** | 2 | 0 | 4 |
| | **Male** | 0 | 0 | 8 |

#### S6 Overlap of constrained genes

Figure S9 extends Figure 3e by showing all overlaps of sets of constrained genes.

#### S7 Expression divergence of gene groups

Figure S11 shows an extended version of Figure 3f demonstrating that the results do not depend on the reference species. Here, two additional species, *D. pseudoobscura* and *D. virilis* were used as reference species from which rest of the species expression divergence and evolutionary time were computed.

Table S3 extends Table 2 utilizing species *D. pseudoobscura* and *D. virilis*, as reference species (in addition to *D. melanogaster*), for computing expression divergence across time. Additionally, another set of Michaelis-Menten curves were fitted, corresponding to the rows labelled as ref = all, considering all points together irrespective of the species used as reference. This demonstrates that the results are not dependent on the reference species.

#### S8 Time taken by EvoGeneX

We ran both versions of OUCH and EvoGeneX on a simulated example of 1000 genes with 4 replicates for comparing the *Sophophora* vs *Drosophila* regimes, on 100 different invocations using R `microbenchmark` package. Figure S16 shows the summary of the time taken in milliseconds for those 100 invocations by each of the two models. Despite the need of estimating an additional parameter and considering larger input data, and more complex relations between them, EvoGeneX is only about 2 fold slower than the version of OUCH that uses average expression of the replicates to infer from 4 replicates. The version of OUCH that uses all replicates takes far much time.

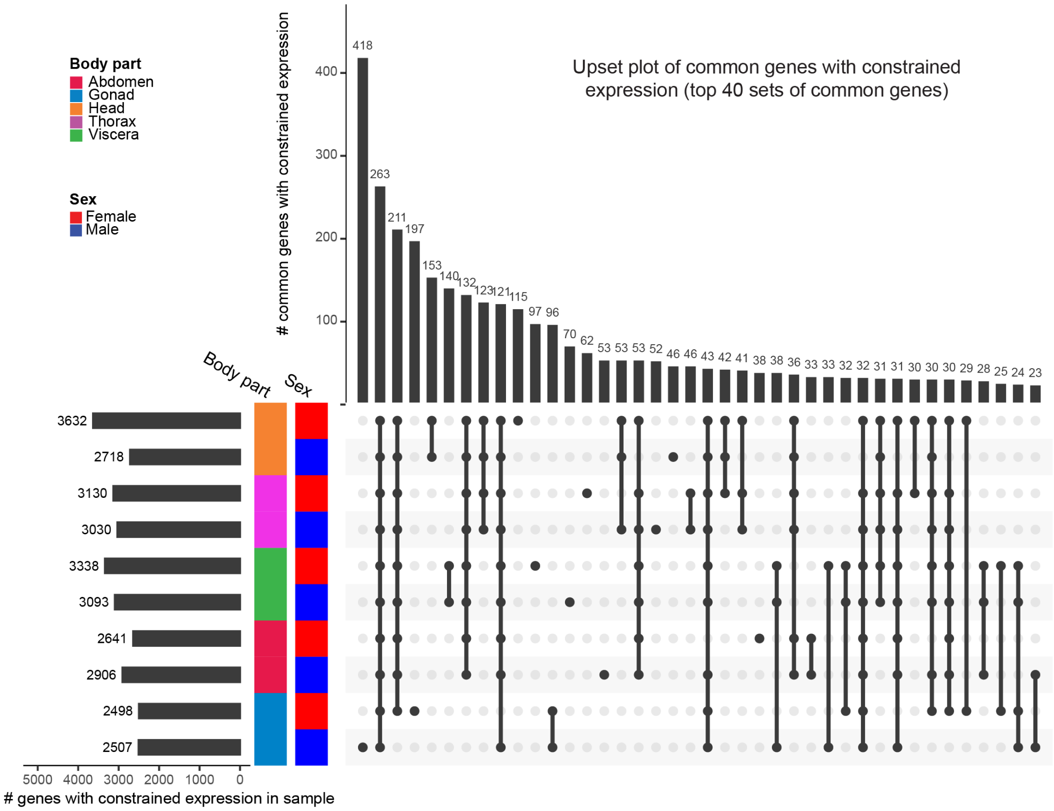


Figure S9: **Overlap of constrained gene groups across tissues and sexes**

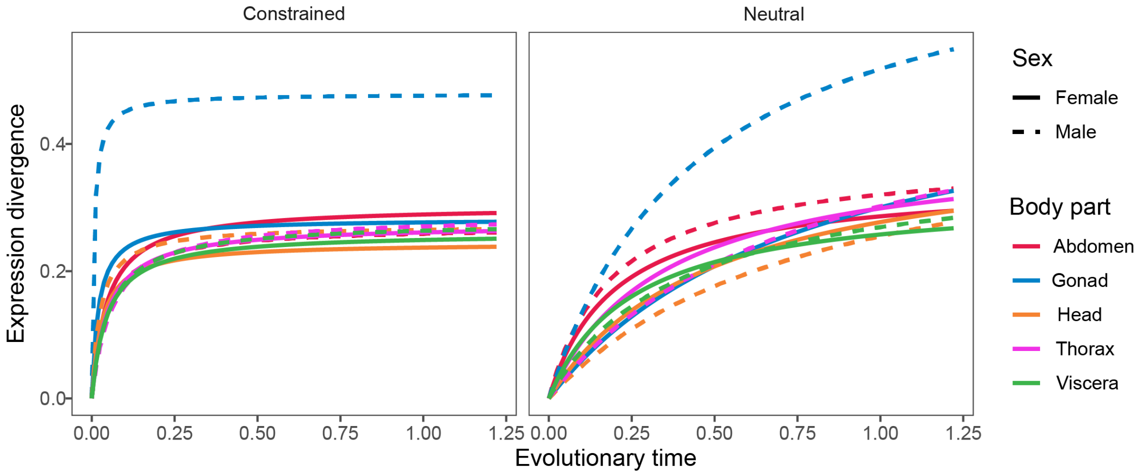


Figure S10: **Michaelis-Menten curves fitted to expression divergence of all species in reference to *D. melanogaster* in all body parts and sexes.**

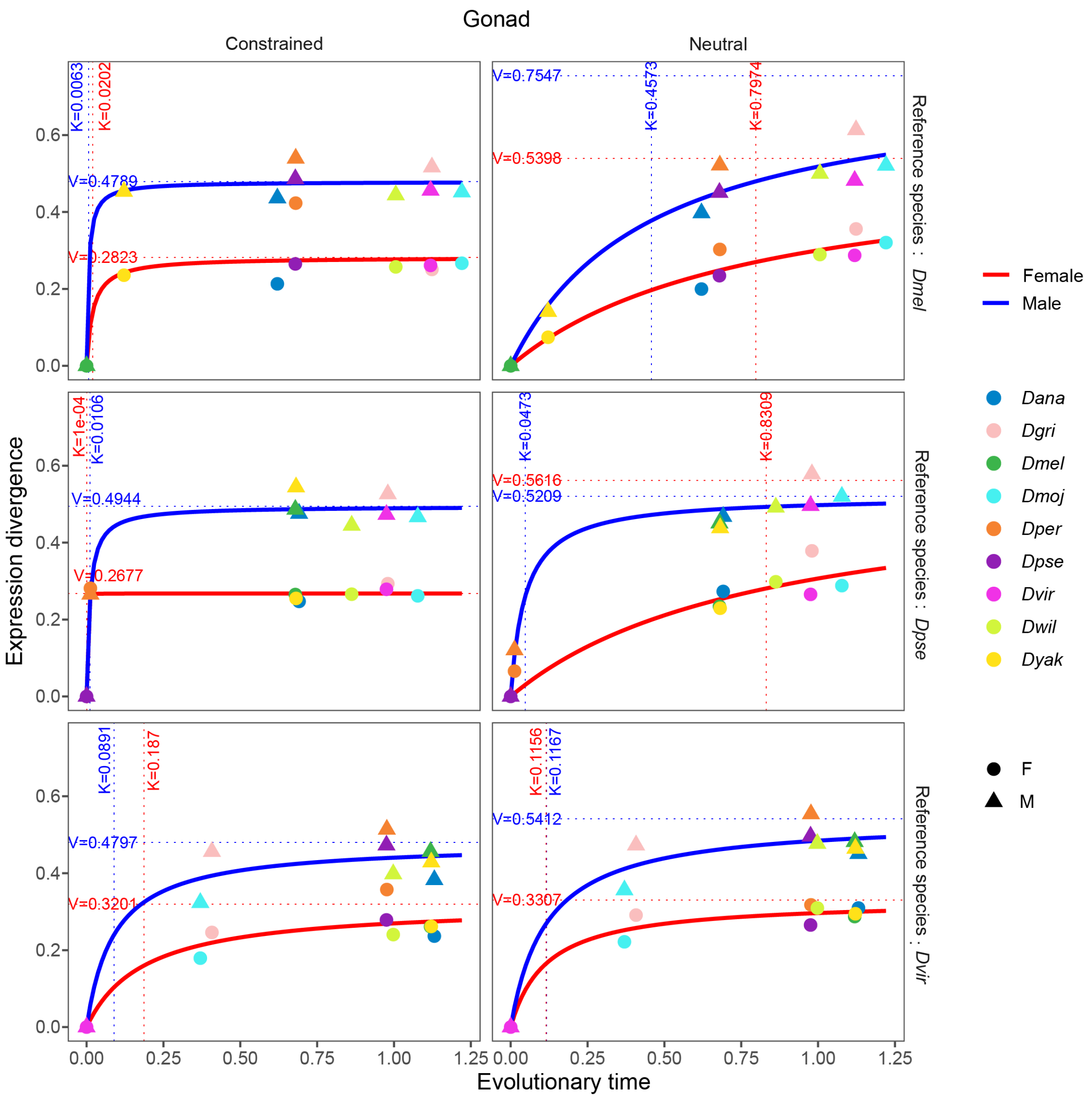


Figure S11: **Expression divergence in gonads over time with respect to three different species,** *D. melanogaster*, *D. pseudoobscura* **and** *D. virilis*, **as reference relative to which the time is calculated.**

Table S3: **Michaelis-Menten curve parameters fitted to expression divergence of different species with respect to three different species (*D. melanogaster*, *D. pseudoobscura*, *D. virilis*) as reference.** The reference 'All' corresponds to case when the divergence points for the three different reference are collectively fitted to a Michaelis-Menten curve.

| | | $V_{max}$ | | | | $K$ | | | |
|---|---|---|---|---|---|---|---|---|---|
| | **Genes** | Constrained | | Neutral | | Constrained | | Neutral | |
| | **Sex** | Female | Male | Female | Male | Female | Male | Female | Male |
| **Ref.** | **Body part** | | | | | | | | |
| *All* | **Head** | 0.221 | 0.244 | 0.336 | 0.330 | 1.211e-04 | 4.931e-04 | 2.472e-01 | 3.169e-01 |
| | **Thorax** | 0.251 | 0.250 | 0.323 | 0.350 | 1.745e-04 | 2.826e-03 | 1.260e-01 | 1.971e-01 |
| | **Viscera** | 0.227 | 0.241 | 0.305 | 0.330 | 1.584e-03 | 3.854e-03 | 1.599e-01 | 2.034e-01 |
| | **Abdomen** | 0.273 | 0.253 | 0.324 | 0.384 | 3.074e-04 | 1.769e-03 | 1.253e-01 | 1.356e-01 |
| | **Gonad** | 0.266 | 0.469 | 0.398 | 0.634 | 1.000e-04 | 9.559e-03 | 3.360e-01 | 2.547e-01 |
| *Dmel* | **Head** | 0.245 | 0.272 | 0.417 | 0.459 | 3.174e-02 | 2.506e-02 | 5.041e-01 | 8.012e-01 |
| | **Thorax** | 0.274 | 0.288 | 0.403 | 0.532 | 4.843e-02 | 6.337e-02 | 3.486e-01 | 7.621e-01 |
| | **Viscera** | 0.261 | 0.278 | 0.323 | 0.377 | 4.665e-02 | 5.348e-02 | 2.537e-01 | 3.996e-01 |
| | **Abdomen** | 0.303 | 0.271 | 0.343 | 0.382 | 4.647e-02 | 4.613e-02 | 1.996e-01 | 1.927e-01 |
| | **Gonad** | 0.282 | 0.479 | 0.540 | 0.755 | 2.022e-02 | 6.266e-03 | 7.974e-01 | 4.573e-01 |
| *Dpse* | **Head** | 0.219 | 0.241 | 0.536 | 0.648 | 1.000e-04 | 2.109e-04 | 9.640e-01 | 1.494e+00 |
| | **Thorax** | 0.252 | 0.248 | 0.784 | 1.555 | 1.000e-04 | 2.162e-03 | 1.631e+00 | 3.918e+00 |
| | **Viscera** | 0.224 | 0.240 | 0.563 | 0.869 | 1.065e-03 | 3.106e-03 | 9.735e-01 | 1.993e+00 |
| | **Abdomen** | 0.274 | 0.261 | 0.281 | 0.346 | 1.000e-04 | 1.769e-03 | 2.966e-02 | 2.217e-02 |
| | **Gonad** | 0.268 | 0.494 | 0.562 | 0.521 | 1.000e-04 | 1.065e-02 | 8.309e-01 | 4.733e-02 |
| *Dvir* | **Head** | 0.223 | 0.243 | 0.293 | 0.277 | 3.879e-02 | 4.096e-02 | 8.298e-02 | 1.024e-01 |
| | **Thorax** | 0.246 | 0.239 | 0.288 | 0.288 | 1.000e-04 | 1.000e-04 | 1.000e-04 | 1.000e-04 |
| | **Viscera** | 0.219 | 0.252 | 0.273 | 0.299 | 1.076e-02 | 9.006e-02 | 4.893e-02 | 5.565e-02 |
| | **Abdomen** | 0.270 | 0.261 | 0.316 | 0.380 | 1.488e-02 | 4.177e-02 | 5.989e-02 | 9.329e-02 |
| | **Gonad** | 0.320 | 0.480 | 0.331 | 0.541 | 1.870e-01 | 8.910e-02 | 1.156e-01 | 1.167e-01 |

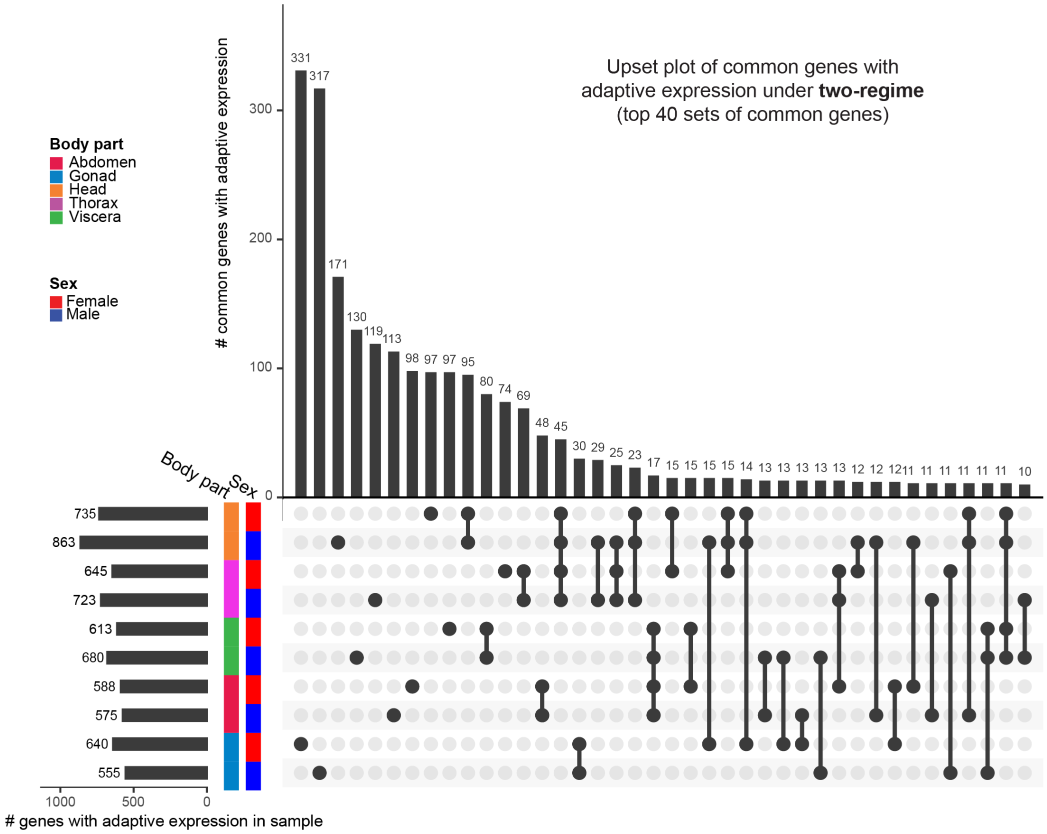


Figure S12: **UpSet plot of top 40 sets of common adaptive genes under two-regime.**

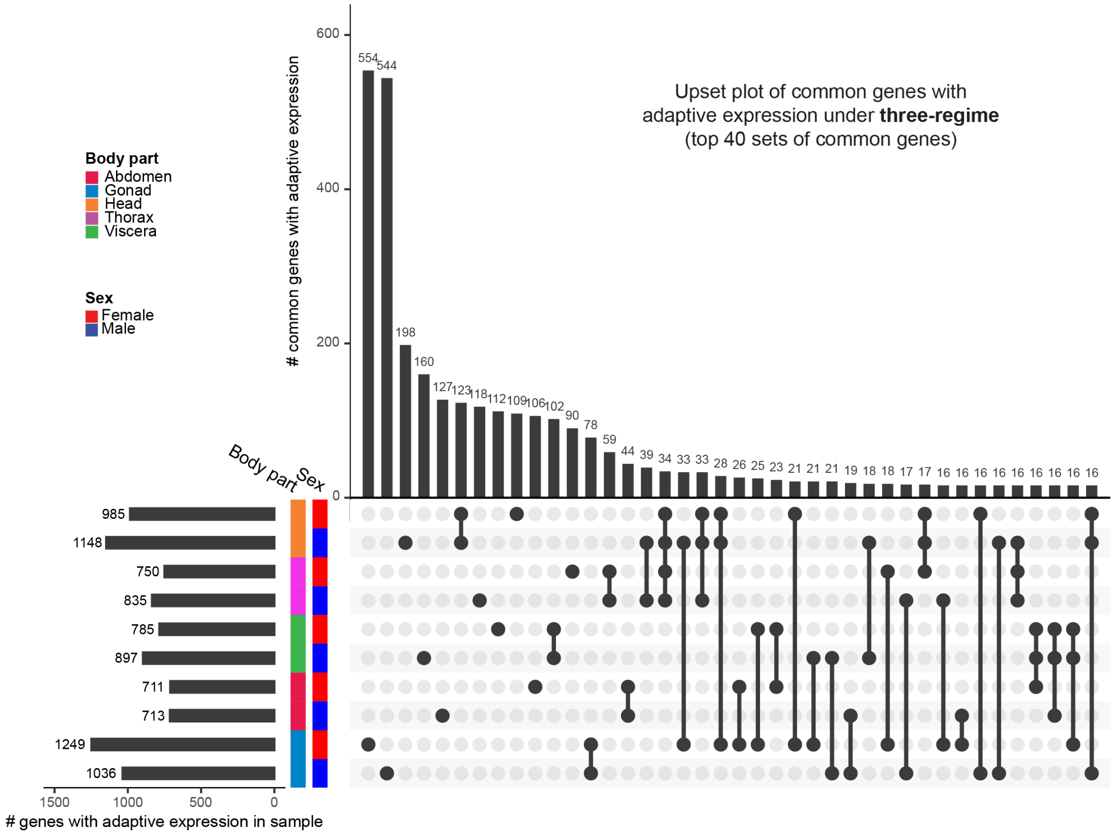


Figure S13: **UpSet plot of top 40 sets of common adaptive genes under three-regime.**

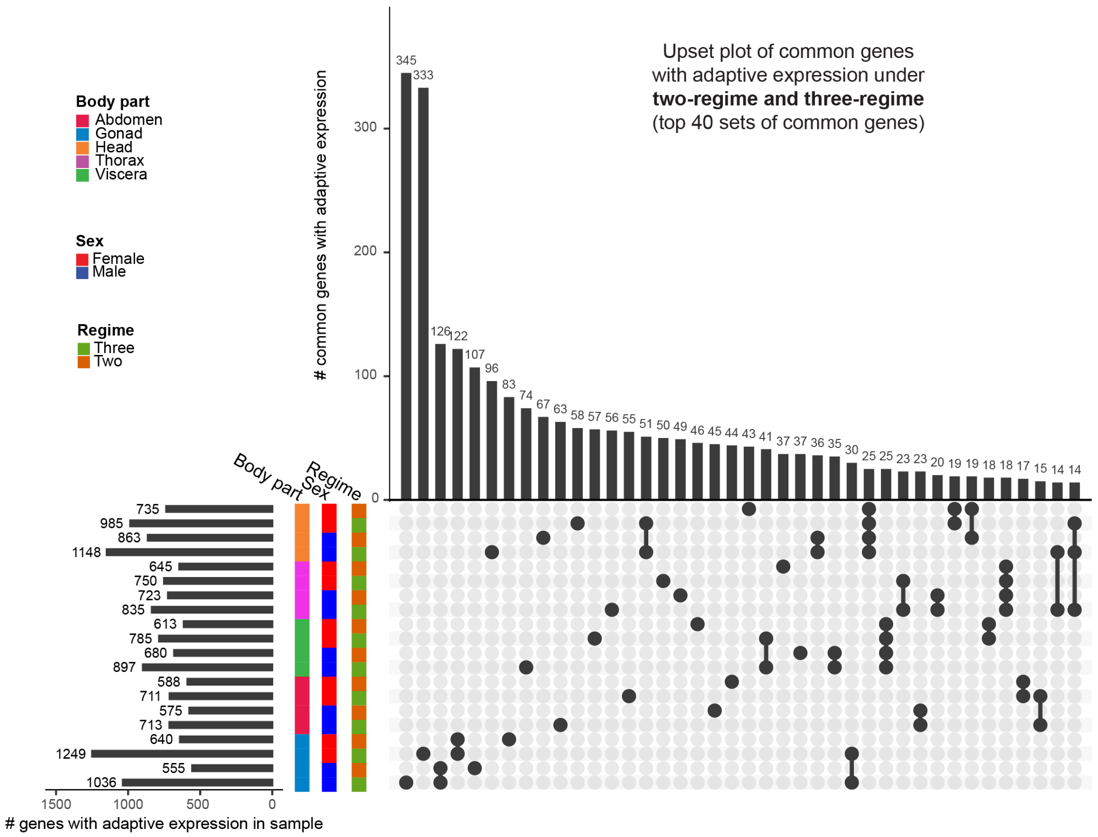


Figure S14: **UpSet plot of top 40 sets of common adaptive genes under both two-regime and three-regime.**

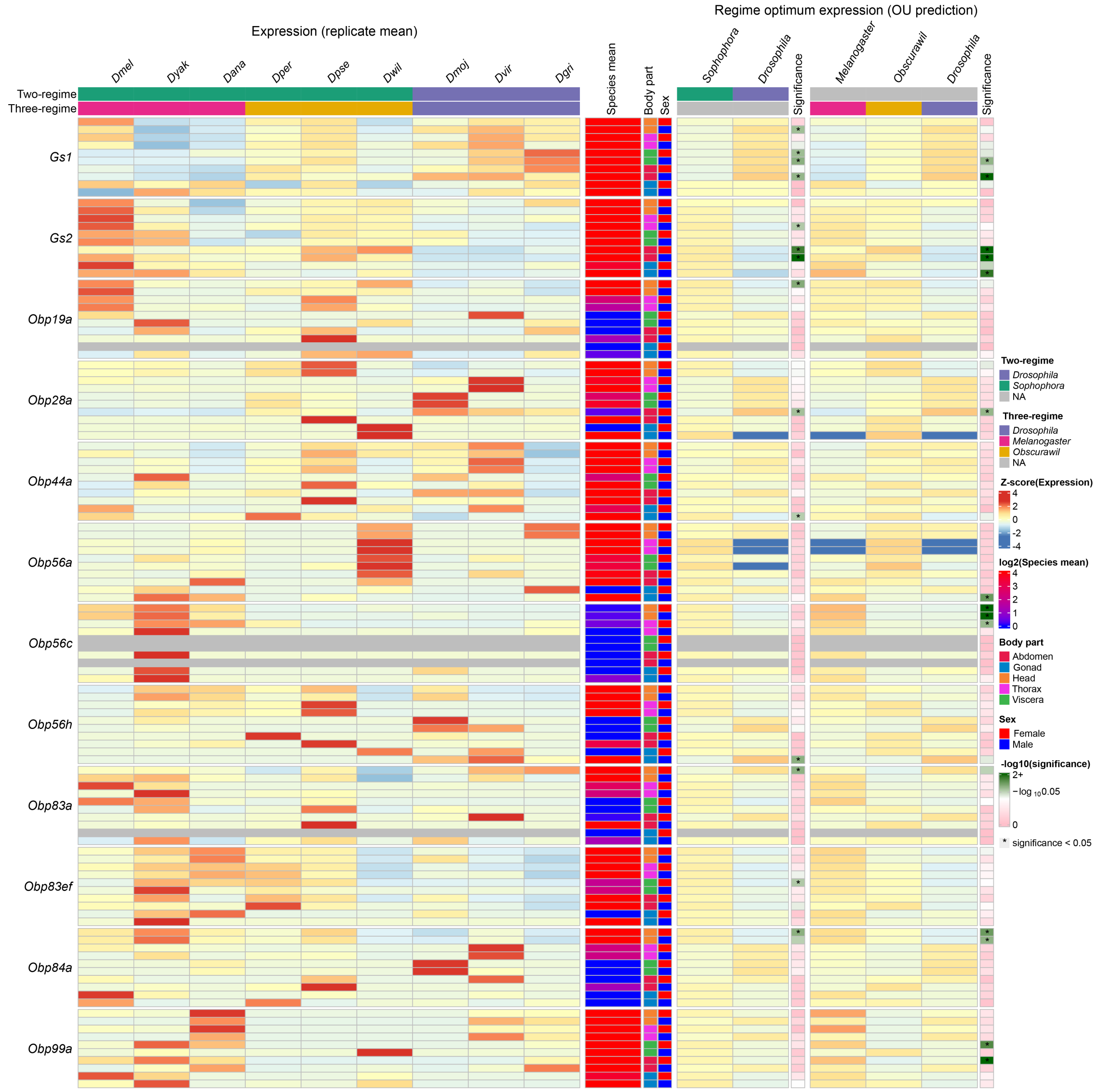


Figure S15: **Detailed view of expression evolution of example adaptive genes in all body parts.**

| Method | Min | Lower quant. | Mean | Median | Upper quant. | Max | Runs |
|---|---|---|---|---|---|---|---|
| **EvoGeneX** | 90 | 93 | 94 | 94 | 95 | 98 | 100 |
| **OUCH.AV** | 44 | 51 | 53 | 53 | 54 | 61 | 100 |
| **OUCH.MV** | 394 | 414 | 424 | 423 | 435 | 458 | 100 |

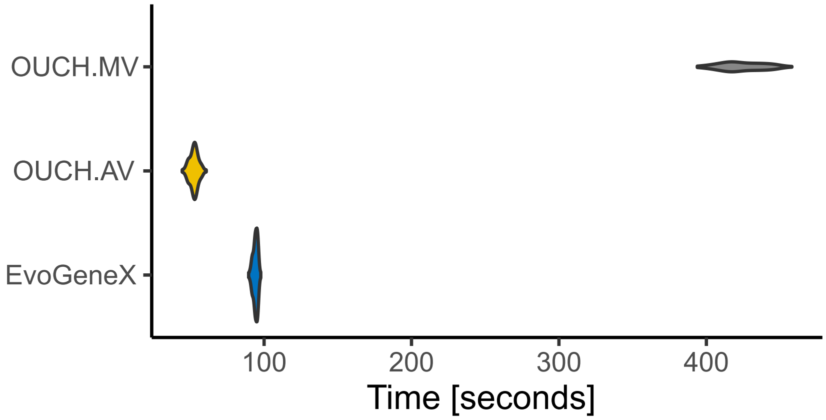


Figure S16: **Time taken in seconds by OUCH and EvoGeneX on an example simulation for two-regime.**
